# Reconsidering the management paradigm of fragmented populations

**DOI:** 10.1101/649129

**Authors:** Oren Kolodny, Michael R. McLaren, Gili Greenbaum, Uma Ramakrishnan, Marcus W. Feldman, Dmitri Petrov, Ryan W. Taylor

**Affiliations:** Department of Biology, Stanford University, USA; Stanford Program for Conservation Genomics, Stanford University, USA; Department of Ecology, Evolution, and Behavior, The Hebrew University of Jerusalem, Israel; Department of Population Health and Pathobiology, North Carolina State University, USA; National Centre for Biological Sciences, Tata Institute of Fundamental Research, Bangalore, India

## Abstract

Habitat fragmentation and population declines call for informed management of many endangered species. The dominant paradigm for such management focuses on avoiding deleterious inbreeding effects in separated populations, by facilitating migration to maintain connectivity between them, an approach epitomized by the “one migrant per generation” rule. We show that this paradigm fails to take into account two important factors. First, it ignores an inherent trade-off: maintaining within-population genetic diversity is at the expense of maintaining global diversity. Migration increases local within-population genetic diversity, but also homogenizes the meta-population, which may lead to erosion of global genetic diversity. Second, this paradigm does not consider that because many fragmented species have declined in numbers only within the last century, they still carry much of the high genetic diversity characteristic of the historically large population. The conservation of a species’ global diversity, crucial for evolutionarily adaptation to ecological challenges such as epidemics, industrial pollutants, or climate change, is paramount to a species’ long-term survival. Here we discuss how consideration of these factors can inform management of fragmented populations and provide a framework to assess the impact on genetic diversity of different management strategies. We also propose two alternative management strategies to replace the “one migrant per generation” dogma.

---

Genetic diversity is considered an important factor in a species’ likelihood of avoiding extinction [1–3]. Genetic diversity can be quantified in different ways; two prominent measures are allelic diversity, i.e. the number of alleles in a population, and heterozygosity, the probability that two gametes sampled at random from a population differ at a certain locus [4]. Genetic diversity, which includes alleles that are under strong selection as well as neutral and even slightly deleterious alleles, encapsulates the evolutionary potential of a population. It serves as a reservoir of potential adaptations to novel environmental and evolutionary challenges [5–9]. Such challenges include new pathogens, climate change, or environmental stress by pollutants. In meta-populations, high diversity, even if not widely distributed between populations, is important. Thus, for example, differential susceptibility of different amphibian populations of the same species to the deadly chytrid fungus may be partially attributable to previously-neutral genetic differences between these populations [3]. Genetic diversity is also important for another reason: low heterozygosity is associated with increased occurrence of deleterious phenotypes due to recessive alleles. Deleterious recessive alleles are common across vertebrates’ genomes, but as long as heterozygosity is high they are usually found in heterozygous individuals that suffer no deleterious effects. Low heterozygosity can thus lead to decreased fitness of many individuals, termed *inbreeding depression*, which in some cases may lead to population decline or extinction [10,11].

Avoidance of inbreeding depression is the focus of many current management plans for endangered species whose populations have been reduced and fragmented. This is often attempted by facilitating connectivity between populations, by translocating individuals between sites or by enabling migration along corridors. It is being considered or implemented for a broad range of species such as tigers [12], rhinos [13], and others [14], and is reflected in the large investment in wildlife corridors across man-made barriers such as highways [15]. Such facilitated migration decreases the frequency of mating between relatives and increases heterozygosity at the local level; however, it also homogenizes the genetics across the connected populations, increases the rate of erosion of genetic diversity at the global scale, and decreases overall evolutionary potential ([3,16–20], see Supplementary sections 1 and 2).

The “one migrant per generation” (OMPG) rule, the epitome of the current management paradigm, was originally proposed to emphasize the role of gene flow in population differentiation, as a balance between local (within-population) and global genetic variation (at the species’ level) [21–23]. It is often interpreted in conservation biology as the “at least OMPG rule”, i.e. a minimal threshold for sufficient gene flow between populations, with many management plans striving to increase inter-population connectivity as much as possible [23]. However, it has been pointed out previously by Templeton and others [3,24] that using the OMPG as a threshold rather than as a theoretical balancing point favors, in practice, maintenance of local heterozygosity over retention of species’ evolutionary potential. These two goals do not always overlap: the latter is dependent on the maintenance of the species’ global diversity, between as well as within populations. Moreover, the OMPG rule assumes that populations are in equilibrium, and does not consider the fact that the populations of species of conservation concern were, just a few decades ago, much larger than they currently are (see, e.g., [23]). The consequence is that a precious legacy of the historically-large populations still exists: much of the genetic variation, and especially allelic diversity, characteristic of large populations, has not yet been lost (Supplementary Sections 1 and 2). The contemporary subsample from those populations represents genetic diversity much greater, sometimes by orders of magnitude, than would be expected from such small populations if genetic diversity had already reached equilibrium. If frequent migration among the remaining populations takes place, genetic diversity will rapidly decrease towards a new low-diversity equilibrium, characteristic of the current small overall population size.

This can be illustrated by considering the diversity at a single genetic locus in a certain species. Assume that 200 years ago this species’ total population was large, say, 1,000,000 individuals. Assuming the species’ population had been fully connected for a long time and was at equilibrium, the expected level of heterozygosity would have been 0.97 and approximately 400 different alleles for this genetic locus would have been expected to be found in the population (the details and calculation of this example are in Supplementary 5). Imagine that following hunting and fragmentation, the species’ global population (*meta-population*) was very recently reduced to 1000 individuals, fragmented into multiple small populations. Because individuals survived the population crash at random, much of the original genetic diversity would still be represented in the smaller contemporary meta-population: heterozygosity would still be expected to be 0.97, and approximately 150 alleles would still expected in the meta-population, representing the legacy of the previous genetic diversity. If frequent facilitated migration between the remaining fragments is carried out, effectively uniting them to form a single population, most alleles would be lost over time. A population’s equilibrium genetic diversity is dependent on its size, and so drift would rapidly eliminate almost all the diversity: the expected equilibrium heterozygosity for a population of 1000 individuals is 0.04, and the expected allelic diversity at equilibrium is below 2. Alternatively, if no migration is carried out, drift would rapidly lead to the fixation of a single allele in each population fragment. However, the changes in allele frequencies due to drift are random and thus many more of the alleles that existed in the original meta-population would avoid overall extinction by stochastically fixing in at least one of the population fragments. If, for example, the species is fragmented to 10 populations, 9 different alleles would be expected to still be represented at equilibrium (see calculation in supplementary 5), and global expected heterozygosity would remain very high: 0.87.

The tension between maintenance of local and global genetic diversity has been pointed out in many studies and relies on well-known theory of population genetics [3,5,16–18,23,24]; however, this has had limited influence on the current paradigm for management of fragmented species. In order to increase the likelihood of fragmented species’ long-term survival, management must consider protection of both local and global genetic diversity. Management should also take into account the meta-population structure of fragmented species and build upon the legacy of precious genetic diversity that can be preserved for the species’ future benefit. In what follows, we present a computational framework, in terms of the number of population fragments maintained and the rate of migration between them, that shows the impact of alternative management plans on genetic diversity as measured by local and global heterozygosity. This framework includes parameters that apply to the endangered species, and is implemented as an interactive tool, available online, for evaluating alternative management scenarios. Further, we propose two simple qualitative management strategies for when and how to carry out migration between populations, facilitating genetic rescue among populations as needed. Finally, we discuss the possible consequences of recurring genetic rescue and the choice of minimal population sizes that would prevent fixation of deleterious alleles of large effect size.

## Modeling the influence of migration on genetic diversity

At the core of our proposed paradigm is the assumption that global genetic diversity is key to long-term survival of species, in agreement with Aldo Leopold’s suggestion that keeping as many of the parts as possible is necessary for “intelligent tinkering” ([25]). To achieve this goal, we use a simple mathematical derivation that computes the changes over time of mean heterozygosity, the most common measure of genetic diversity used in conservation [11,26], in the meta-population and within populations. We can explore scenarios that differ in the size and number of populations and in the rate of migration between them (the model is presented in Supplementary 2; we use ‘heterozygosity’ as defined above, known also as ‘expected heterozygosity’ [4]). Because many species are characterized by a ‘historical legacy’ of high genetic diversity compared to the equilibrium state for the current meta-population size, and because much of the diversity that needs to be protected is selectively neutral or near-neutral, especially in currently small populations, our model focuses on the impact of random drift on neutral genetic diversity under different management scenarios.

Figure 1 depicts the dynamics of genetic diversity, measured as heterozygosity, under one management scheme in which fragments are fully connected via migration and another with no migration at all, and compares them to the dynamics under an OMPG management scheme. It illustrates what is expected to happen to global heterozygosity and to within-population heterozygosity under each. The no-migration scheme is expected to preserve global heterozygosity at much higher levels than both of the other schemes. The OMPG scheme, although commonly thought to retain local genetic uniqueness of populations (e.g., [23]), genetically homogenizes the separate populations at a rate similar to the fully-connected management scheme, and drives loss of global diversity at a similar expected rate. However, it maintains higher local diversity within populations than the no-migration scheme. Figure 1 demonstrates the tradeoff between retaining local and global diversity. We suggest that management plans should not follow a specific one-rate-fits-all migration rule, such as OMPG, but should aim to balance global and local diversity: Global genetic diversity should be maintained at the highest levels possible, retaining as much as possible of the remaining historical genetic variation. At the same time local genetic diversity must be maintained at a level that is necessary to avoid population-level inbreeding depression. Our model, also implemented as an online tool, allows the exploration of the effect on local and global heterozygosity to be expected with different migration rates and numbers of populations.

**Figure 1:**
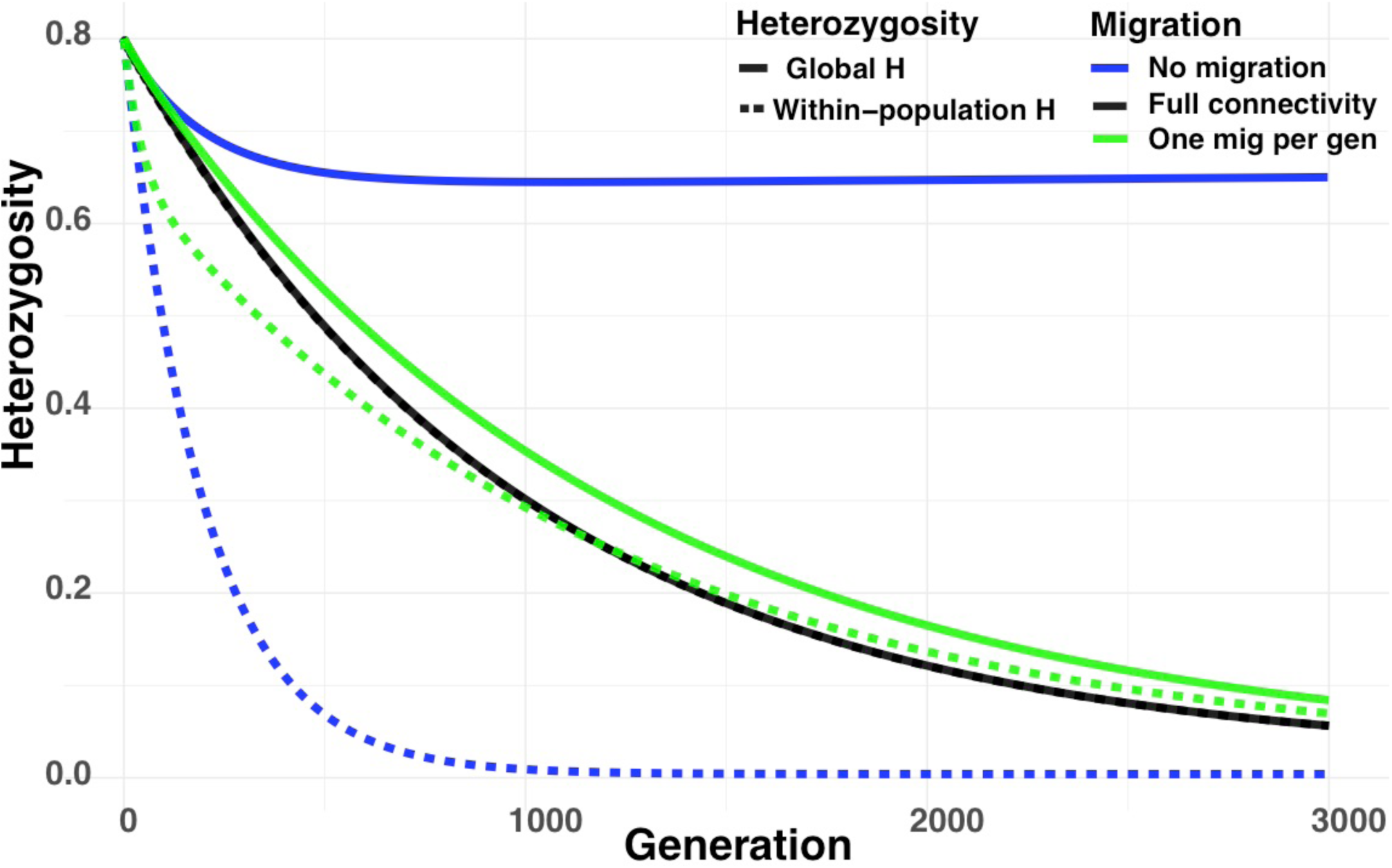
Expected global and local heterozygosity under different regular migration schemes. A species whose historical meta-population size was *N*_*e*_ = 100,000, and was reduced to 5 fragmented populations, each characterized by *N*_*e*_ = 100. Maintenance of full connectivity (black) leads to an identical decrease in global and local heterozygosity. Complete avoidance of migration leads to retention of much of the original genetic diversity, reflected in high global heterozygosity (solid blue), but also to rapid decline in heterozygosity within local populations (dashed blue). Migration at a rate of one migrant per generation leads to coupled decline of global and local heterozygosity (solid and dashed green, respectively), that diverges only slightly from that under a scheme of full connectivity. Assumed rate of mutation is 10^-5^, similar to rates found in microsatellites among vertebrates. Examples for dynamics in populations with different parameters are found in supplementary 1. Results are numerical calculations based on equations 3 and 6 in Supplementary Section 2, and were produced using our computational tool, available at https://ryantaylor.shinyapps.io/gmfp/.

## Proposed management schemes

The management paradigm we propose does not include pre-defined regular rates of assisted migration between populations. Instead, we suggest that migration should be facilitated only if population-level inbreeding depression becomes imminent. The impact of inbreeding and the level of heterozygosity at which it becomes a problem are unclear, particularly in light of cases in which populations persist despite having been founded by very few individuals or having undergone severe bottlenecks (e.g. [27,28]). Among studies that show inbreeding effects in the wild, the vast majority show decreased fitness of inbred individuals but do not address whether inbreeding has an impact at the population level (e.g. [29,30]). However, the population, not the individual, is what matters for species’ conservation; the reason for preventing inbreeding should be to avoid local population declines in a way that could lead to local extinction and the concomitant loss of population-specific genetic diversity (see discussion in supplementary 2). We suggest that management should avoid regular migration of individuals among populations; instead, we suggest that managers should ‘play the (genetic) cards’ sparingly, and make use of inter-population genetic diversity in order to prevent local extinctions only when necessary. We propose that the way to do this is by employing ‘genetic rescue’ ([31,32]) – migration of individuals from one population to another – only if the receiving population shows signs of decrease in population size that are suspected to be related to lack of genetic diversity (*scheme 1*). Figure 2 illustrates recurring application of genetic rescue into our model (see also Supplementary 2). We find that even under a broad range of conservative assumptions, this scheme is superior to regular migration of individuals between populations in the spirit of the OMPG rule. It leads to a slower rate of loss of global heterozygosity (Figure 2, solid orange curve), while maintaining viable local populations. Supplementary 1 discusses which populations to use as sources for migrants in genetic rescue.

**Figure 2:**
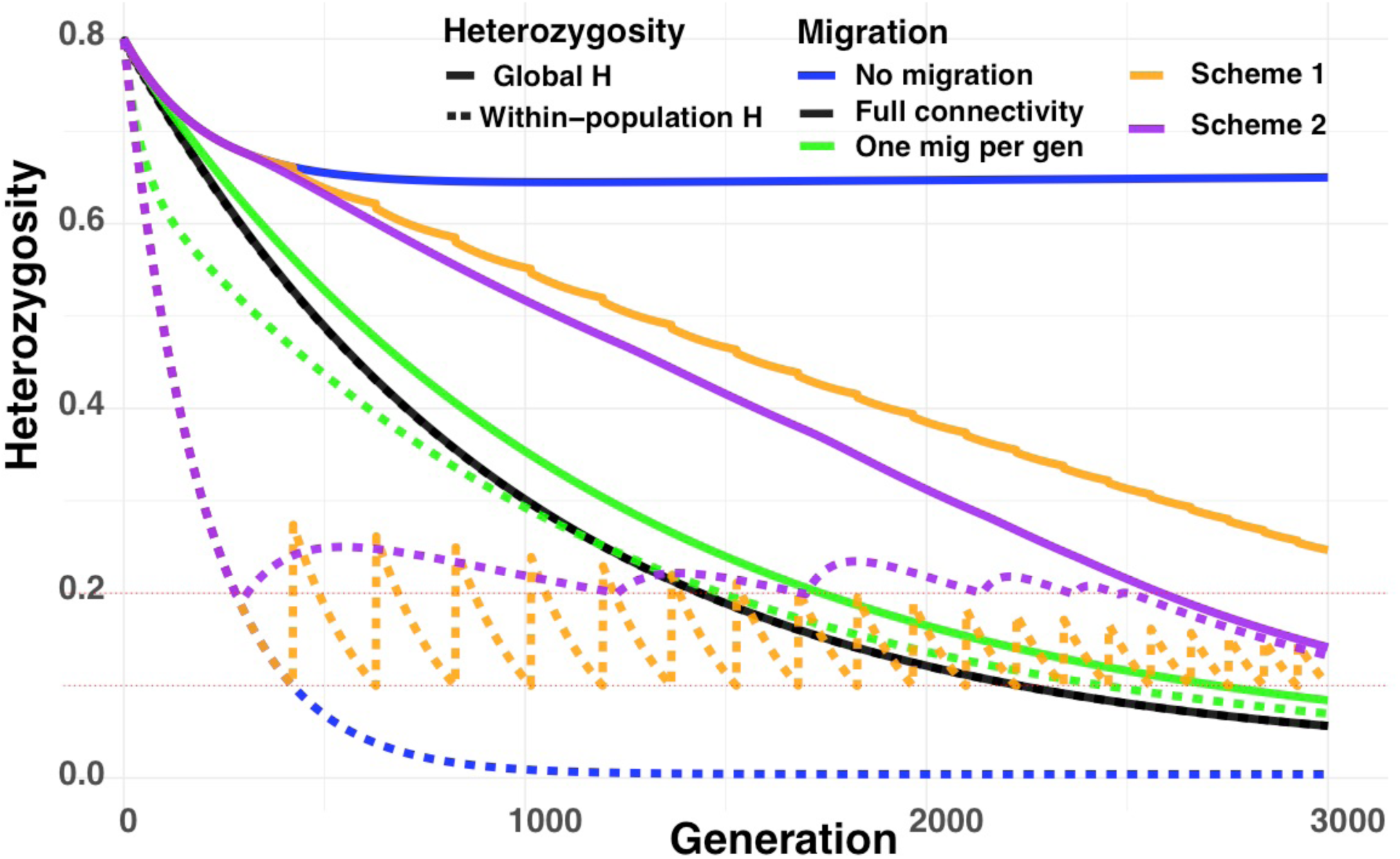
Proposed management schemes and comparison to the OMPG. The primary scheme (scheme 1) includes no migration, except when a population begins to decline, in which case genetic rescue via migrants from another population is facilitated. Using a derivation based on the pseudo-hitchhiking model [33] (Supplementary Section 2), we assume that genetic rescue leads to replacement of the variant found at each genomic site in a certain fraction of the individuals in the rescued population, *R=*0.2 in this illustration, by the allele for that site that was carried by the rescuing individual. The figure depicts the expected global and local heterozygosity (solid and dashed orange, respectively) assuming for this illustration that such rescues need to be implemented whenever local heterozygosity reaches *H*=0.1 (merely for the sake of illustration, since rescues are not expected to be required in a regular manner, and are not expected to be directly related to heterozygosity). The alternative scheme we propose (scheme 2, purple) is the facilitation of minimal migration while avoiding a pre-defined level of heterozygosity that is deemed critical with respect to the risk of inbreeding depression (here, *H*_*critical*_=0.2): no migration until this threshold is reached, followed by a shift to a regular rate of migration which is updated every few decades as necessary (rates chosen for this illustration are, respectively: no migration, one migrant per 20 generations, one per 10 generations, one per 6 generations, one per 4 generations, and one per 2 generations; details of this illustration, and illustrations for other parameter values, are in supplementary 1). The population sizes are as in Figure 1; the plots for the regular migration schemes, identical to those shown in Figure 1, are presented for comparison (black, green, blue). Scheme 1 allows maximal retention of global diversity for the longest period of time, avoiding *H*=0.2 for thousands of generations longer than does OMPG, while maintaining viable local populations. Scheme 2 allows the retention of less global diversity than scheme 1, but may be more appropriate in some cases (see text), and also far outperforms OMPG both in maintenance of global diversity and in the duration for which it allows the avoidance of the critical threshold of local heterozygosity (purple and green curves; avoidance of the threshold for hundreds of generations longer than the OMPG). The figure was produced using our online tool and can be further explored at: https://ryantaylor.shinyapps.io/gmfp/.

Although evidence suggests that genetic rescue can be very rapid, there may be species in which demographic dynamics make this less likely to succeed or too risky due, for example, to stochastic fluctuations in population sizes. There are also populations whose close monitoring is impossible, and for these population decline may go unnoticed until very late. For such cases, we propose an alternative management scheme (*scheme 2*), which can be viewed as an informed version of the OMPG (details in supplementary 1). In scheme 2, managers define the critical level of heterozygosity that they believe must be avoided in order to prevent population-level inbreeding depression. Initially, no migration is carried out, thus preventing unnecessary loss of global diversity. When a population’s heterozygosity nears the chosen threshold, migration at a low rate is facilitated, maintaining heterozygosity above the threshold. This rate of migration is updated every few decades to maintain the population above the critical heterozygosity threshold for as long as possible. This scheme outperforms all schemes of regular migration between populations, both in the length of time that it allows avoidance of the critical threshold of heterozygosity in each population (Figure 2, dashed purple), and in the global diversity that is retained (Figure 2, solid purple; also Supplementary 1).

A potential challenge in maintaining small populations is the avoidance of fixation of deleterious mutations: efficiency of purifying selection against deleterious mutations is dependent on population size. The detailed dynamics of such purging in small populations are debated ([34–36]), and the relative importance of deleterious mutations in the conservation context is unclear. We propose that an advisable precaution is to determine the size of the managed populations in a way that is conservative in its likelihood of preventing fixation of deleterious mutations of large effect. Using standard assumptions and denoting this effect size by *s* and the *effective population size* by *N*_*e*_, deleterious mutations that satisfy *N*_*e*_ *• s* >> 1 have a negligible probability of fixing. For example, if each population had *N*_*e*_*=*40 or more, then *N*_*e*_ *• s* > 2 for deleterious mutations that reduce fitness by 5%, thus effectively preventing their fixation (see supplementary 4 for details and discussion of relation between *N*_*e*_ and census population size). Accordingly, where local populations are very small, we advise the creation of a regional structure, such that each *region* consists of a number of populations that together meet the requirement for the minimal *N*_*e*_ and constitutes a management unit. Within each region, migration of individuals should be facilitated such that the populations will be effectively fully connected or nearly so. Among regions, migration will be facilitated only when genetic rescue is required.

## Summary

The paradigm shift we propose for management of fragmented species, with the goal of increasing the likelihood of long-term species’ survival, is based on a simple model of the dynamics of genetic variation. We challenge the currently-prevalent paradigm, which is focused on increasing inter-population connectivity and does not consider the historical legacy of high genetic diversity, a valuable asset for species long-term survival. It is based on models that do not directly address genetic diversity and the risk of inbreeding depression, but focus on population divergence, a measure that is frequently irrelevant to this risk.

Beyond the scope of the genetic diversity model that we focus on here lie a number of additional important genetics-related arguments against facilitated migration between populations that we do not discuss. These include the risks of outbreeding depression and the loss of local adaptations. Consideration of these factors lends further support to the approach we are suggesting.

Our focus here has been on genetic diversity, which is only one factor among many in species’ conservation. Our study does not address demographic or ecological considerations, which may often pose a greater risk for species’ survival in the short term (e.g., [37,38]). Such considerations include the risk of Allee effects, risk of extinction due to stochastic fluctuations in population sizes, and risk of spreading disease and parasites between populations by migration. Finally, removing direct threats to the species’ existence such as hunting and road kill, and maintaining or restoring ecosystem and habitat integrity and resilience, are additional crucial steps.

As conservation efforts are forced to shift towards management of discrete protected areas and their resident populations, informed strategies should be adopted that both maintain high global genetic diversity and avoid local population extinctions. Our paradigm suggests that to do so successfully, populations should be closely monitored but that active maintenance of connectivity be reduced to the bare minimum necessary, recognizing that the one migrant per generation rule may be, in many cases, one migrant too many.

## Software

We offer an online tool for the assessment of different migration rates on global and local heterozygosity, as shown in Figure 1 and 2. The code and link can be found at https://github.com/rwtaylor/GMFP.

## Supporting information

Main Text

## Acknowledgements

We thank Elizabeth Hadly, Stephen Palumbi, Jonathan Pritchard, Oded Berger-Tal, Roee Maor, and Shirley Bar-David for helpful comments. O.K. is supported by the John Templeton Fund. This research was also supported by the Stanford Center for Computational, Evolutionary, and Human Genomics, and the Morrison Institute for Population and Resource Studies at Stanford.

