## Supplementary material for "Reconsidering the management paradigm of fragmented populations": Main Text

#### Content

Supplementary 1: Management schemes and examples of model predictions for illustrative examples

Supplementary 2: Mathematical derivation of change over time in local and global heterozygosity in a structured population

Supplementary 3: Review and discussion of related studies and test cases

Supplementary 4: A brief summary of population structure theory

Supplementary 5: Supporting calculation for the illustrative example in the main text

**Supplementary 1: Management schemes and examples of model predictions for illustrative examples**

The two main components of the conservation paradigm that we propose are (1) Consideration of the trade-off between retention of high within-population diversity and the retention of global diversity, and (2) Appreciation of the fact that many of the fragmented species still retain the precious historical legacy of high diversity that characterized the historically large population. Different management schemes can be devised that take these two factors into account. We advocate the use of a management scheme that completely avoids assisted migration among populations as long as populations are viable, showing no signs of decline. However, in some cases it may be advisable to adopt a different scheme, for example if intensive monitoring of populations is not possible and there is a risk that population declines would go unnoticed for long periods of time, or if the species' life history suggests that it would not be amenable to effective genetic rescue if necessary.

In the following we (1) outline two management schemes that take into account the two primary components of our paradigm[1], (2) discuss additional considerations and practical heuristics that may be useful for decision making, and (3) use our modeling framework to analyze a number of "toy examples" for which we predict the outcomes on different time scales of some of the proposed management schemes with realistic parameter values.

[1] A few previous studies, most prominently those of Caballero and co-authors, have proposed valuable conservation principles and tools that highlight similar aspects, albeit with an emphasis or suggested management strategy that differs from ours [8–11,42,44–47]. A major difference between our approach and the approach advocated in many of these studies is that their focus is on controlled mating between particular individuals with consideration of their individual genetics, while we propose a low-intervention scheme that controls only the migration dynamics between populations. The approaches are complementary, and the appropriateness of each depends on factors such as population sizes, species' life history, and the available funds and technological resources available for the management program.

**Management schemes and management considerations**

**Foreword: Deme sizes**

Our analysis focuses on genetic diversity and its possible influence on population-level dynamics in the context of providing evolutionary potential for adaptation and of avoidance of inbreeding depression. Another consideration is the risk of fixation or of increase in prevalence of deleterious alleles. As described in the main text, we propose to address this by creating a regional structure, such that each *region* consists of a number of populations that together meet the requirement for the minimal $N_e$ and constitutes a 'management unit' ('region' is used to refer to these management units, and not as a geographic locality). Within each region, migration of individuals will be facilitated such that the populations in it will be effectively fully connected or nearly so. The following management schemes are described in terms of the rules of facilitated migration between regions (see also heuristic approach for region size determination, below).

Our study does not address demographic considerations. Such considerations may, depending on the ecology of the species of interest, play a prominent role in determining region sizes and their management. If populations within regions are very small, such that stochastic fluctuations may lead to locally-extreme sex biases or to extinction, demographic considerations may dominate. These would dictate the maintenance of very high connectivity within each region to prevent local extinctions from occurring (see, e.g., [1]).

In what follows, for simplicity, the term 'population' is used as shorthand for 'the whole population of a certain region', since the small populations within each region are effectively fully connected).

**Management scheme 1: *Waste not, want not* – no regular migration, use genetic rescue as required according to population monitoring**

Predicting populations' genetic dynamics to a high resolution, even under neutral genetic assumptions, is notoriously difficult, particularly in small populations. Moreover, it is nearly impossible for wild populations, as it requires many assumptions about the species' life history and details about the particular population of interest. Although an attempt at such prediction is recommended for informed management, we suggest avoiding reliance on such model predictions to prescribe preemptive migration. Instead, we suggest relying on empirical population-specific evidence to determine whether migration is required. In this way, global genetic diversity – which is best retained by keeping regions separate – is never 'wasted': migration is kept to a necessary minimum.

We suggest that facilitated migration should be carried out if at least one of two conditions is met:

(1) The region's population is in numerical decline for reasons that are likely due to genetics (i.e. due to poaching or habitat destruction, for example, genetic rescue is irrelevant). In these cases, the decline may be due to inbreeding depression or to a lack of adaptive alleles or allele combinations that may overcome ecological challenges such as a changing climate or a novel pathogen. Genetic rescue may ameliorate both conditions.

(2) The population reaches a state in which most individuals have crossed a critical threshold of individual heterozygosity to an extent that puts the population as a whole at risk. In many species, individual-level fitness declines precipitously beyond a certain threshold of inbreeding coefficients (see supplementary 3). If such a "natural critical value" exists for what is likely to be a catastrophic reduction in population viability, it is advisable to avoid reaching a state in which a large majority of individuals in the population are beyond that threshold. Importantly, empirical evidence suggests that no absolute value for such a threshold exists that would apply to all species [2–4]. This empirical critical value is species-specific, and may even be population-specific due to differences between populations in their history (e.g. [5]). In the absence of a natural heterozygosity threshold, we suggest reliance on condition (1) only.

The primary implementation challenge in this management scheme, and to a large extent in any scheme that wishes to balance the trade-off between maintaining global and local genetic diversity, is the assessment of the population's state. This assessment requires close monitoring of population demography, and would also benefit from some longitudinal genetic sampling to assess individual-level inbreeding coefficients via homozygosity. Such monitoring, however, is necessary for any informed management that transcends the simplistic rules of thumb that follow a "one rate fits all" approach. Such a step is timely and necessary, and is possible today more than in the past, thanks to technological developments that allow efficient genetic and demographic monitoring.

An important special case may arise if for some reason implementing a regional structure is inapplicable, and some populations are maintained at very small sizes, on the order of less than 20 breeding individuals (e.g. [6,7]). In such cases, it may be possible to track the pedigrees of all (or most) individuals in the population, and their coefficient of inbreeding can be calculated. If the population reaches a state in which most individuals are inbred to an extent analogous to mating between half-siblings, we suggest that the extremely small population size combined with high inbreeding should prescribe the implementation of genetic rescue even in the absence of clear evidence for genetically-associated population decline.

**Management scheme 2: *The informed X-migrants-per-generation rule* – model-based determination of migration rates and their timing**

Management scheme 2 is an informed version of the OMPG paradigm. It considers the trade-off between global and local diversity, and although it prioritizes – as does the OMPG – complete avoidance of a certain threshold of local heterozygosity to avoid risk of inbreeding depression, and places retention of global diversity as a secondary consideration, it optimizes the management plan not to "overshoot" this goal. It aims at avoiding migration as much as possible, allowing the heterozygosity within regions to decrease to a pre-defined critical threshold, and then it maintains the population at that heterozygosity threshold with recurrent migration. The rate of migration is very gradually increased over time, so as to constantly be near the minimal rate that is necessary to maintain the population above the critical threshold.

We suggest that management scheme 1 should be preferred whenever possible, as it relies on the specific condition in the population and is founded on the actual "ground truth" of the necessity of facilitating gene flow, i.e. it uses measures that relate directly to the goal being optimized – the population's viability, tested via monitoring its growth or decline. Management scheme 2 focuses not on population viability directly, but on a secondary proxy: the risk that may be associated with population-level inbreeding depression. This focus is problematic, because it is *a priori* unclear when inbreeding depression begins to have a significant effect at the population level, how this effect quantitatively relates to extinction risk, and how severe this increased risk is compared to the cost of mixing populations and risking loss of more genetic diversity than is absolutely necessary. However, such an approach may be appropriate in some cases.

As in the OMPG and related schemes, the premise of management scheme 2 is that population viability is likely to be significantly reduced once the majority of the individuals in it cross a certain threshold of individual heterozygosity, as discussed in the previous section. As in the OMPG, the determination of migration rates in this scheme relies on a (necessarily simplistic) model of the dynamics of heterozygosity over time in relation to migration rates between regions, and is less reliant on regular high-resolution monitoring of the population. However, the OMPG is designed to optimize a measure that is not context-specific and is removed from the goal of avoidance of a certain extent of inbreeding: it aims to reach an $F_{st}$, a measure of population divergence, to be 0.2 at equilibrium, a balance point that is not necessarily informative regarding inbreeding or inbreeding depression, particularly in populations outside of equilibrium of genetic diversity. Management scheme 2, instead, relies directly on heterozygosity, which is more informative about inbreeding.

The idea behind management scheme 2 is that – using the derivation that we provide in supplementary 2 and the online interactive tool that we have set up to explore alternative schemes (see below), alongside other potential resources – one can predict – regrettably, with a large degree of uncertainty – the level of heterozygosity in the population as a function of time for different migration schedules. Management scheme 2 proposes to avoid migration between regions as long as they are above a critical threshold of heterozygosity that was chosen to be avoided (this threshold can include a buffer zone where heterozygosity is above the value at which a critical drop in viability is expected). If, or when, a within-region heterozygosity decreases to this threshold, migration at a regular rate between regions is implemented. This migration rate is chosen to maintain the regions' expected within-region heterozygosity above the threshold for as long as possible. This rate is updated once in a while as necessary (see the examples below; for a broad range of realistic parameters, rate updates need to be carried out as little as once in 100 generations). The timing of the switching point between no migration to a regular rate of migration can be pre-defined based on model predictions, or determined empirically (bringing it closer in nature to management scheme 1). Regardless, since model predictions regarding change in heterozygosity are very imprecise, it is highly advisable to monitor the change in heterozygosity in the population over time, and update the rate of migration accordingly.

This scheme also differs from the OMPG paradigm in that it suggests that a species-specific threshold of heterozygosity be set, and it considers the historical inheritance of high genetic diversity: the OMPG is designed for populations in genetic diversity equilibrium, ignoring the fact that most species of interest are far from their equilibrium diversity given their current population size.

Interestingly, management schemes 1 and 2 may converge: conservation plans often set a time horizon, which is defined as the goal for the plan. Such time horizons are typically on the order of 200-500 years (following an assumption that at this time point, or long before, the species' status and management plan will be re-assessed and revised). In some species it might be that, thanks to the historical inheritance of high genetic diversity from the historically large population before fragmentation, the heterozygosity within regions is high enough that even in the complete absence of migration between them, the per-region heterozygosity is not expected to reach the critical threshold within the defined time horizon.

The opposite may also occur: it may be that the current (and expected near-term) overall population size is so small, that even full admixture of all the remaining individuals would not slow down the loss of diversity to an extent that would maintain the heterozygosity above the desired threshold. In such cases a different management strategy should be considered. One option is, in the spirit of management scheme 1, not to assume that a certain heterozygosity exists but to allow individual populations to reach very low levels of heterozygosity, and use genetic rescue between populations when necessary, thus maintaining the species' evolutionary potential, its global diversity, for as long as possible. An alternative, which is preferable if possible, is to engage in individual-level determination of mating events in the population, in order to optimize the retention of global diversity and decrease co-ancestry of mated individuals (see, e.g., [8]).

#### Region size determination

In the main text we outlined a simple approach to calculating an effective population size that is recommended per region in order to prevent the likelihood of fixation of deleterious mutations of large effect. However, linking census population size and effective population sizes is hard and requires many assumptions about the species life history and the specific population. Additionally, it is unclear which deleterious effect size should be chosen as the critical threshold to be avoided and how efficiently selection acts in practice in holding such deleterious alleles at bay.

In the spirit of management scheme 1, we propose an heuristic for determining regional structure (definition of management units) that attempts to avoid population admixture as much as possible, in order to protect global diversity which may be needed for the species to adapt to future evolutionary challenges. We propose to initially choose intentionally small region sizes, i.e. to set out by defining regions such that they include fewer populations than might in practice be required to meet the necessary effective population size. Over time, if these initial regions' populations show signs of decline, we suggest following scheme 1 of applying genetic rescue, doing so with migrants from neighboring regions. If a region requires a second and third genetic rescue, we suggest attempting such rescues from the same region of origin as the first attempt. If one or both of the regions continues to decline, the two regions should be fully admixed to compose a single region.

This strategy will allow the gradual convergence towards the region sizes necessary to maintain viable populations, without "overshooting" in a way that would be wasteful of global genetic diversity. Notably, it is reliant on close monitoring of the population in each region.

#### Choice of sources for genetic rescue

The primary goal of our proposed conservation paradigm is the maintenance of genetic diversity at the species level, in order to retain the species' evolutionary potential to cope with future environmental challenges. Obviously, the mere retention of diversity is not enough: a clear strategy of its utilization when required is also necessary. We propose to do so in the form of genetic rescue in response to populations' needs, as signaled by population demographic decline.

The effective implementation of facilitated migration, either for genetic rescue or as part of a migration scheme at a regular rate, is a significant challenge, that has been studied and discussed extensively (e.g. [6,9–11]), giving rise to valuable insights and implementation schemes. Here we highlight a few possible strategies and considerations that focus on the choice of source population from which to conduct facilitated migration between regions when needed.

**Strategy 1 – Hierarchical meta-regional structure:** In this approach, neighboring regions would be used for genetic rescue, according to pre-defined clusters of regions, in a way that gradually creates a hierarchical tree-like structure among all regions that are managed. In this tree-like structure, the closer two regions are on the tree, the more likely they are to have been used for genetic rescue of one another. At the extremes, after a number of genetic rescues had been carried out between a pair of neighboring regions, as outlined in the previous subsection, this pair could be viewed as one region and regular frequent migration between the populations that compose it should be facilitated. This strategy slows down the meta-population admixture compared to a scheme in which the source for genetic rescue is chosen arbitrarily every time such rescue is needed, and it thus retains as much global diversity as possible for as long as possible.

Although it is conceptually and practically simpler to use neighboring regions for recurring genetic rescue and eventual unification if needed, the same scheme can be applied independently of geographical distance between regions. The hierarchical structure that is used to define and describe which regions are closer to one another does not need to be aligned with geographic structure of the population. Arguments in favor of a structure that aligns with geographical distance include its simplicity, a reduced likelihood of transmission of disease that could otherwise not spread, a reduced likelihood of leading to outbreeding depression, and the emergence of an isolation-by-distance gradient which is characteristic of natural populations, is robust to rare events of non-facilitated migration between populations, and aligns with the possibility that in the future reclaiming of land for habitat restoration or protection would allow population growth and reconnect fragmented populations. Arguments in favor of a structure that is incongruent with geographical distances are that, because the original populations were in practice rarely fully admixed as our model assumes, such a structure increases the likelihood that the migrants for genetic rescue would have significantly different genotypes from those in the receiving region, increasing both the likelihood of alleviating inbreeding depression and providing novel genetic adaptations.

Notably, the hierarchical structure that is created according to the strategy described above, which describes the emerging genetic relation between regions, can either be pre-defined in full, or allowed to arise as a result of consecutive decisions regarding sources for genetic rescues as the need for these arises. Earlier decisions, in this case, constrain the range of choices for later decisions, but the full hierarchical structure does not need to be defined ahead of time, and flexibility should be retained in order to choose sources optimally with respect to the state of affairs in each population at the time of the rescue's execution and to additional considerations (see the following strategy; the two strategies are not mutually exclusive).

**Strategy 2 – Evolutionary rescue approach:** The assumption behind this strategy is that the genetic rescue is necessary, at least in part, because the receiving population's habitat undergoes an environmental change and the population lacks allelic variants that may help to address this challenge. The idea is that such alleles may be found in another population whose historical environmental conditions were (or still are) similar to the novel environment of the receiving population. Such cases may occur as a result of climate change, that renders a north-latitude habitat similar to that previously existing in southern latitudes, for example, or the emergence of a pathogen in a certain region that had previously been known (and successfully dealt with, at least to some extent) in a different region. The source of individuals for genetic rescue can thus be chosen based on each population's historical conditions. This strategy and the previous one, that proposes a hierarchical regional structure for genetic rescue and according relatedness between populations, are not mutually exclusive.

We note that these suggestions leave open many aspects of genetic rescue, such as whether it is preferable to attempt genetic rescue multiple times between the same two regions, or whether it is preferable to choose every time a different pair of regions from the same cluster in the (possible pre-defined) hierarchical structure. Additionally, the decisions in each situation would be greatly dependent on the specific case, including, for example, the hypothesized reason that underlies the necessity for a particular instance of genetic rescue, the specific population and the meta-population's structure, differences in environment, differing risks of inbreeding and disease transmission in migration between different pairs of regions, and many other factors. Many valuable suggestions are found in studies that have been done of the topic (e.g. [9,12–14]), and we expect that understanding of the involved mechanisms and the relevant considerations will increase as experience with the implementation of genetic rescue accumulates.

**Long term species conservation**

In the long term, the ultimate solution for a species stable existence, from the perspective of conservation genetics, is an increase in its populations' sizes. This scenario may be possible for some species, particularly those whose population crash occurred due to stressors that can, or have been, removed. This is the case of the Asiatic wild ass that has been re-introduced in Israel, for example: from a founding population of 11 individuals, the population has by now increased to a few hundred, and continues to grow [15,16]. However, for most species significant population growth in the future seems unlikely.

The conservation strategy that we propose, which emphasizes the retention of species-level diversity, serves both scenarios well. For species that are going through a bottleneck and whose population will grow in the future, retention of diversity through maintaining populations as separate fragments will allow much of the historical diversity to be retained; once population growth occurs, the risk of global loss of diversity due to drift will be reduced even if all the populations are allowed to reconnect. The proposed strategy may save genetic diversity that would otherwise require many centuries to be regained. The majority of species, whose populations will remain small, are expected to face a deluge of environmental challenges in the near future (some due to human-induced changes, such as climate change and prevalence of pollutants). For these species, the retention of global diversity – its species-level evolutionary potential – alongside its timely use for populations' survival in the form of genetic rescue when such rescue is needed, is crucial for survival.

**Is there a threshold of critical heterozygosity?**

Both scheme 1 and scheme 2 require assessment of whether a critical heterozygosity threshold exists and what that threshold is – either for regular monitoring and as a trigger for action, as in condition 2 of management scheme 1, or for pre-determination of a long-term target of minimal heterozygosity that regional populations are allowed to reach before facilitated migration is implemented. Assessing such heterozygosity theoretically requires many assumptions, and thus may yield dubious conclusions. We suggest that a preferable approach is to glean insight about heterozygosity-fitness relations from empirical data. Ideally, data from the species of interest or from a species close to it is best (even though differences may exist in the relation between the inbreeding coefficient ($F$) and fitness, even between populations of the same species, depending – for example – on differences in historical population sizes, bottlenecks, and purging dynamics).

We point out a number of findings from the literature that may serve as points of reference. More than anything, they demonstrate the extent to which the relationship between $F$ and fitness, particularly population-level fitness, is unclear. A discussion of inbreeding is found in supplementary 3. Although whole-genome sequencing is becoming available at costs that are affordable enough for it to be used in regular monitoring of species of conservation concern, we focus in the current discussion of heterozygosity values inferred from microsatellites, as the majority of studies in the literature, and most implemented conservation plans, still employ this measure.

Large well-mixed populations of intermediate- and large-bodied vertebrates are characterized by *expected heterozygosity* [17] values in the range of 0.25 to 0.8 (e.g. [18–20]). Bottlenecked small populations exhibit a broad range of heterozygosity values, ranging from those whose heterozygosity does not significantly differ from that of larger populations, to values as low as 0.05 ([18,20,21]). Interestingly, as discussed in supplementary 3, some studies find that populations with low heterozygosity exhibit reduced performance on various fitness-related measures such as mean clutch size or mean body size [21], while in others there is no effect, or an effect only on particular measures of fitness but not on others [22]. Moreover, in some cases even populations that exhibit reduced performance in some fitness related measures, such as island populations with low heterozygosity suffering from an increased parasite load, also exhibit high performance in other measures related to population viability compared to their mainland counterparts, assessed – for example – via population density ([23]). Importantly, even populations that have low heterozygosity and reduced values in measures that act as proxies for population-level fitness do not necessarily fare worse than others, and some such populations seem to have existed stably at small population sizes for hundreds of generations ([18,20,21,24]; see further discussion in supplementary 3).

For the sake of the illustration of our model and comparison of alternative management schemes, we assume that the critical threshold of heterozygosity is 0.2, and we assume a mutation rate of $10^{-5}$.

**Toy examples: characteristics for considerations and comparison of management outcomes**

To explore the time scales and relative rates of change in genetic diversity under different management schemes, we apply our model to three toy examples that differ in their original population sizes and in the extent to which their populations have been reduced: (1) a species whose effective population size[2] was reduced 100-fold from 500,000 to 5000, (2) a species whose overall effective population size was on the order of 100,000 and has been reduced 200-fold to 500, and (3) a species whose overall population size was on the order of 100,000 and has been reduced 1000-fold to 100. The parameter values of toy example (1), henceforth *Toy1*, are on the order of magnitude assessed for many intermediate-sized mammals, and also near a lower bound regarding both historical and present population sizes for many others, that are targets of local and global conservation efforts such as the population of European badgers in north-west Europe ([25]), the population of Sika deer in japan ([26]), the populations of mountain Gazelle subspecies in the east Mediterranean and the Levant [27], and others [28]. The parameter values of toy example 2, *Toy2*, are on the order of magnitude assessed for some large predators such as tigers [29]. Toy example 3, *Toy3*, represents an even more extreme reduction in the population size of the current population compared to the historical one, to an exceptionally small current size, and may perhaps best represent a species whose populations were reduced due to habitat destruction and has remained only in very few specific sites, such as some endemic island species [30], some freshwater fish [31], and amphibians [32,33].

[2] Henceforth in this section, *population size* refers to variance effective population size; the relation between this and the census population is discussed in supplementary 4.

Our derivation, and accordingly the illustrations below, does not assume any particular strategy for choice of migrants or choice of genetic rescue: it assumes that the source of these, whenever facilitated migration is carried out, is chosen at random from among all regions. Informed choice of the source of migrants increases the likelihood of its success and/or the retention of global diversity, as discussed above, so in this sense our illustrations are conservative in assessing the advantages of the schemes we propose over the OMPG and similar strategies.

Wherever rates of migration or rescue are assumed, they refer to rates of effective migrants, i.e. a rate of one migrant per generation means that one migrant successfully migrated and joined the regional gene pool. In some instances this may require a number of migrants, and the choice of these migrants as well as the way in which migration is carried out – e.g. in terms of sex, age, seasonal timing, etc. – are important. These have been the focus of much study, reviewed, for example, in [12,14,34–37].

**Example 1: Intermediate current population size (*Toy1*)**

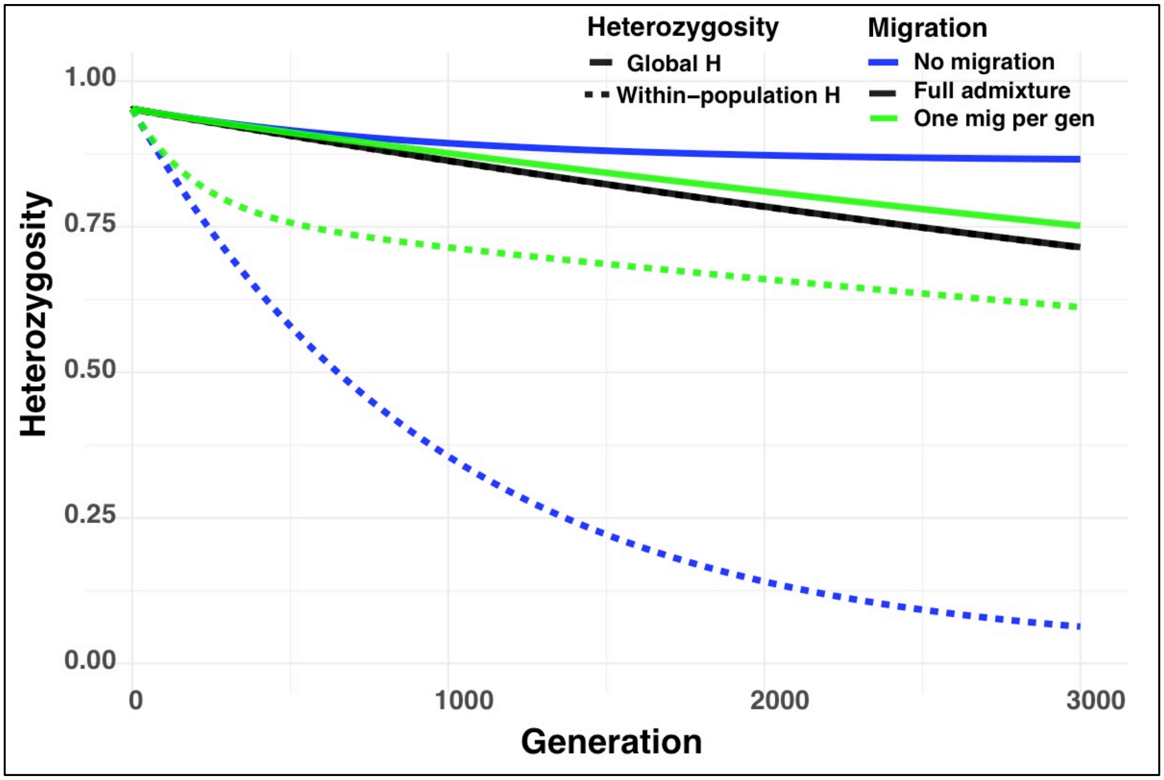


**Figure S1.1:** Expected change in heterozygosity over time in a species of 5000 individuals, divided into 10 populations, which historically numbered 500,000 individuals. Mutation rate: $10^{-5}$. Curves depict heterozygosity change under management schemes of no migration (blue), one migrant per generation (green), and full admixture (black).

The parameters of this example (*Toy1*; see figure S1.1 caption) are characteristic of many intermediate sized vertebrates whose conservation status as species is as yet of only limited concern thanks to their relatively large population sizes. Management with respect to connectivity of such species is typically not via facilitated migration of particular individuals, but through efforts to create corridors in the landscape that allow animals to move between fragments. Such efforts include road and railroad animal crossings (underpasses and bridges), for example.

The characteristic time horizon of conservation plans is 100-500 years. On this time scale, in Toy1 (even for an annual species), even a strategy of no migration at all does not lead to local heterozygosity that is near the assumed critical threshold of $H$=0.2. On this timescale, global heterozygosity is affected to a limited extent, regardless of the management strategy, i.e. even full admixture leads to only limited reduction in heterozygosity, because the timescale on which drift occurs is dependent on the population size, which is fairly large. However, significant differences in the extent to which global diversity is retained arise at intermediate timescales, and are quite salient at 1000 or 1500 generations into the future. This suggests that for species characterized by parameters similar to these, it is preferable to avoid between-region migration, thus retaining as much global diversity as possible while still not compromising the local populations with respect to risk of inbreeding. Moreover, the budgets invested in maintaining such between-region connectivity, which are often quite significant (discussed in , e.g., [38,39]), would be better used for other conservation goals.

**Example 2: Small overall population**

**A**

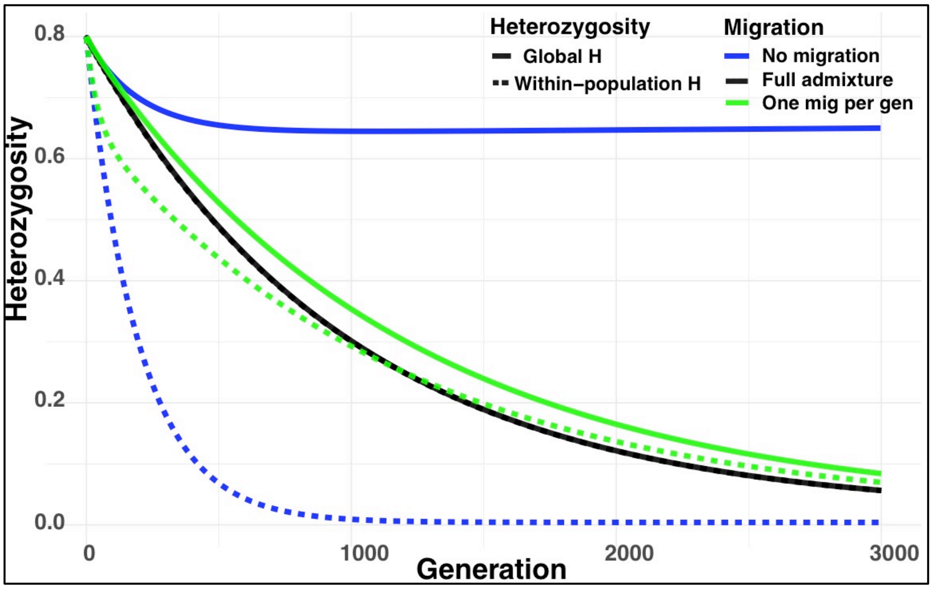


**B**

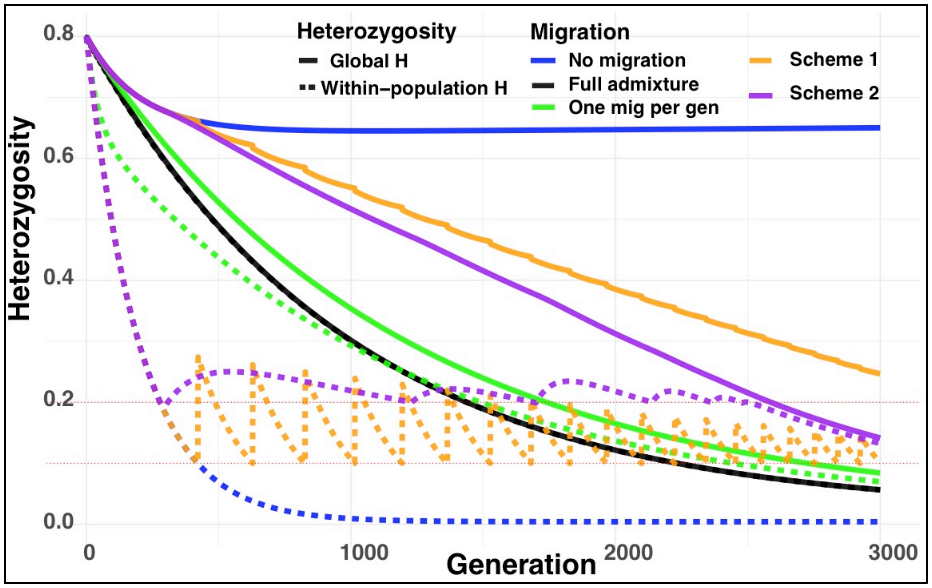


**Figure S1.2:** Expected change of heterozygosity over time in a species of 500 individuals, divided to 5 populations, which historically numbered 100,000 individuals. (A) Change under management schemes of full admixture, one migrant per generation, and no migration. (B) Change under the management schemes of (A), compared to change under management schemes 1 and 2 suggested in this study. Mutation rate: $10^{-5}$.

*Toy2* is the example illustrated in the main text (Figures 1 and 2). It is characteristic of many populations that are actively managed and closely monitored, such as Tigers ([29]; assuming that small fragments are connected to comprise regions with population size on the order of a few dozens or more as discussed in the main text). Here, a no-migration strategy would lead to severe reduction in local heterozygosity, even within the time horizon of typical conservation plans or near it, while the OMPG would not. This might seem to suggest that OMPG is a beneficial strategy in this case. However, figure S1.2b demonstrates that the schemes that we propose in this study far out-perform the OMPG both in the retention of global diversity and in the time span in which they manage to maintain the levels of local heterozygosity above the thresholds that are assumed to be critical.

**Genetic rescue in management scheme 1**: we assume (see supplementary 2) that when genetic rescue is facilitated, as opposed to regular migration, the incoming individuals enjoy a fitness benefit, either because the population they are reaching is suffering from population-level inbreeding depression, which is decreased in the incoming individuals' offspring, or because they carry adaptive alleles that had been missing in the target population. In these cases, some of the migrants' alleles would rapidly sweep to high frequency or fixation. The resulting sweep(s) will lead to the loss of linked neutral variation through the process of genetic hitchhiking. The strength of hitchhiking is determined by a parameter $R$, which is the fraction of gametes in the rescued deme that are replaced by copies of the hitchhiking allele: the neutral allele that is linked to the sweeping allele. In the illustration above, R = 0.2. The genetic variation in the replaced gametes is lost, so that hitchhiking tends to result in a drop in heterozygosity and in allelic diversity within the deme and in the meta-population as a whole. This contrasts with a migrant that arrives in a population as a part of a regular-migration scheme (scheme 2), and has no particular advantage. Such a migration event might thus bring about loss of allelic diversity indirectly, via introduction of an allele that could drift to high frequency or fixation, but it does not induce selective sweeps and sudden loss of diversity.

For the sake of the illustration, we assumed that population decline occurs at $H$=0.1, necessitating genetic rescue (orange, management scheme 1), and that every time genetic rescue is carried out, a proportion R=0.2 of the gametes is replaced by copies of the hitchhiking allele. This leads to a gradually increasing rate of genetic rescues that are required (Figure S1.2b). In reality, it is unclear whether the conditions requiring genetic rescue are likely to be due directly to low heterozygosity, and if they are, how low heterozygosity would have to be in order to necessitate genetic rescue. We suggest that a threshold of 0.1, chosen here merely for the sake of illustration, is quite conservative. On the other hand, it seems likely that R may take on significantly larger values than 0.2 (with R=1 meaning that all alleles in the region were replaced by those of the migrants in the genetic rescue, which is – from a genetic perspective – analogous to an extinction-recolonization event). Management scheme 1 may be superior to management scheme 2 or not, depending on these unknowns that are particular to each species, and even to each specific population. Additionally, since this analysis is of the dynamics of the expected mean heterozygosity, realized dynamics are expected to be much more variable. In the graphs presented above, rescue was initiated every time local heterozygosity reached a value near 0.1, leading to a rescue rate that was initially once every 200 generations, and that increased gradually to every few dozen generations.

**Migration in management scheme 2:** Following the period of no migration, when local heterozygosity approaches the pre-defined threshold, migration is carried out at a certain rate. This rate can be constantly adjusted and migration carried out in a way that would maintain heterozygosity exactly at the chosen threshold. However, both in reality and for the illustration, it is simpler to define a certain rate of regular migration for a long period of time, and increase this rate in a discrete way every time that the heterozygosity approaches the threshold again. In Toy2, migration was switched initially from zero to one migrant per 20 generation, to one migrant per 10 generations, one per 6 generations, one per 4 generations, and finally one every 2 generations, at which point both global and local heterozygosity dropped below the threshold. Notably, this occurs many hundreds of generations after the time point at which this threshold is crossed under the one migrant per generation management scheme. Supplementary 2 provides an alternative, analytically-based scheme, for choosing the periodically-updated rates of migration.

Because the time horizon of major changes in heterozygosity in Toy2 is near that of conservation plans, it may be sensitive to the details of the parameters, e.g. a 2-fold decrease in the size of regions would shift the expected dynamics to be well within the time horizon of many conservation plans. However, the qualitative results hold: under the assumptions of our derivation (e.g., that genetic rescue and facilitated migration can be effectively carried out and influence genetic diversity, as measured by the expected mean heterozygosity) the management schemes that we propose and that consider both global and local diversity outperform any strategy of regular migration, for any given time horizon and choice of critical heterozygosity threshold. They are – by their definition – the least wasteful of global diversity in meeting the chosen goal (ameliorating population declines or avoiding of a certain heterozygosity threshold), thus also increasing the time span during which local heterozygosity can be maintained above a given threshold, because global and local heterozygosity – if any kind of migration is facilitated – are coupled.

**Example 3: A very small population at present**

A

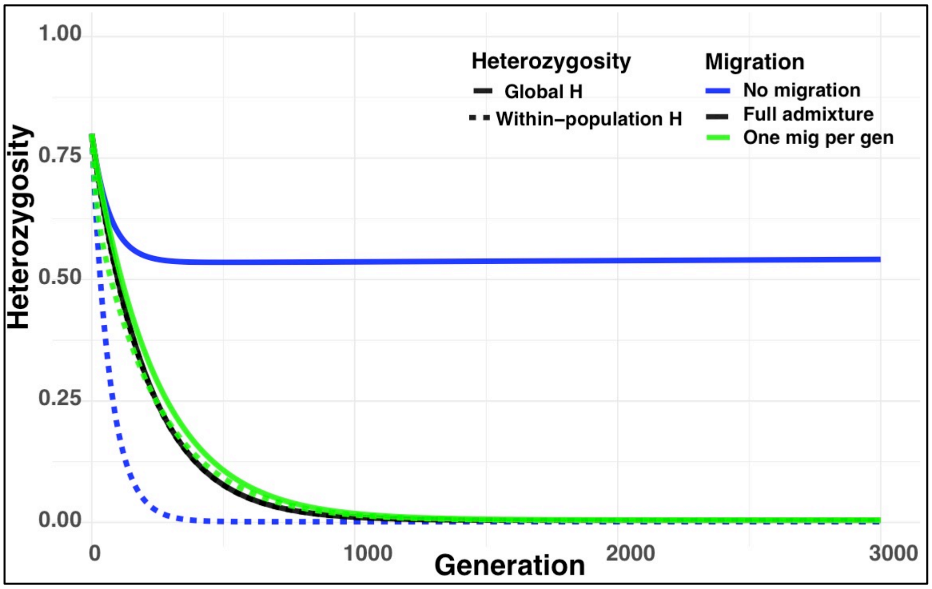


B

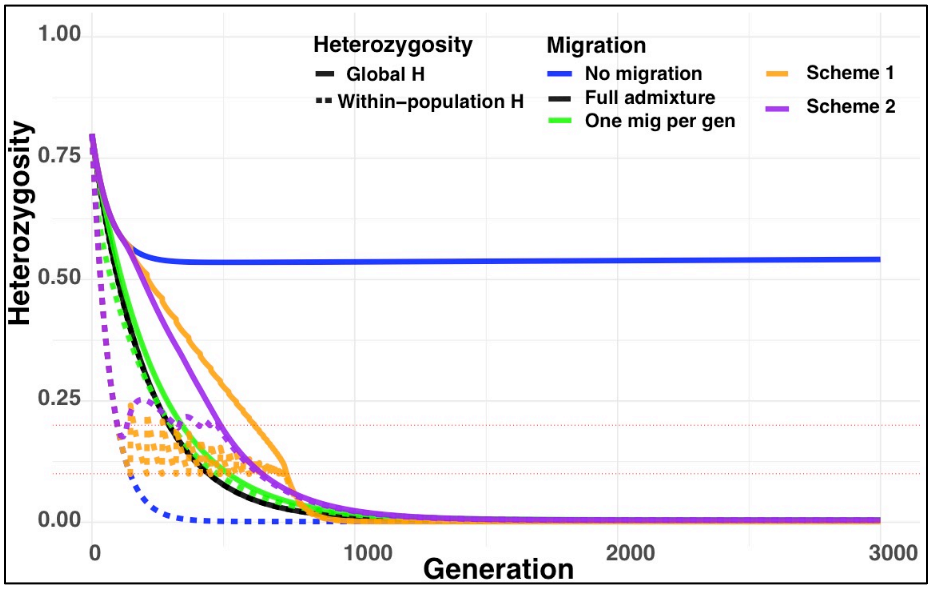


**Figure S1.3:** Expected change of heterozygosity over time in a species of 105 individuals, divided to 3 populations, which historically numbered 100,000 individuals. (A) Change under management schemes of full admixture, one migrant per generation, and no migration. (B) Change under the management schemes of (A), compared to change under management schemes 1 and 2 suggested in this study.

This example (*Toy3*) is similar to the previous one. It differs in that here, the overall population size is so small, that nearing or crossing the critical heterozygosity threshold within the time horizon of many conservation plans is inevitable. Some might intuitively suggest that in such settings there is nothing to do but to mix all the remaining populations (in full, or using an OMPG rule) and hope for the best. The illustration above suggests otherwise: even in cases as in this toy example, an advantage to both global and to local heterozygosity is gained by using migration sparingly and not beyond the minimum required, following either of the schemes proposed in this study, postponing the crossing of the defined heterozygosity thresholds by a few hundred generations (Figure S1.3b; scheme 1, 2: orange and purple, respectively). In particular, within the first 200 generations, which is the primary target-timeline of many conservation programs, the proposed schemes 1 and 2 maintain the populations above the defined critical thresholds (H = 0.2, 0.1 respectively) while retaining global heterozygosity which is higher by 35% or more than the heterozygosity retained under a one migrant per generation scheme.

Notably, since the time scale on which diversity is lost in this case is short, it may be that some species whose populations crashed in the recent two centuries have already lost much of their diversity. Although this is not the case in the example in Figure S1.3, it may sometimes happen that following either of the two schemes we propose would dictate the implementation of genetic rescue (scheme 1) or of migration due to low heterozygosity (scheme 2) at a rate that is similar to an OMPG-style plan, in which case the two paradigms converge. However, such convergence should not be assumed a-priori.

In such small populations it may be feasible and advisable to implement a high-intensity management plan, in which mating between pairs of individuals are manipulated, pairing specific individuals with one another, in a way that maximizes the retention of genetic diversity and increases the effective population size. A number of such endeavors have been successful (see discussions in, e.g. [40–43]), and computational frameworks that allow such optimization are available (e.g. [9]).

### Supplementary 2: Mathematical derivation of change over time in local and global heterozygosity in a structured population

We use mathematical models to study how different migration management schemes affect the dynamics of heterozygosity within individual populations and in the meta-population as a whole. We first analyze the classic finite island model, which illustrates the conflict between the maintenance of local and global heterozygosity and allows us to calculate the expected trajectories of heterozygosity under management plans that include no migration, full admixture, regular rates of migration, or periodically-updated rates of migration (as we propose in management scheme 2). In the finite island model, maintenance of global heterozygosity is always favored by reducing migration; indeed, eliminating migration entirely allows for almost all of the initial diversity to be kept, as illustrated in the main text and calculations in Supplementary 5. The high levels of global diversity achieved under a no-migration policy may be unrealistic if such a strategy necessitates periodic genetic rescues. Populations experiencing genetic rescue will tend to lose neutral genetic variation due to the process of genetic hitchhiking [1, 2]. Thus rescues will result in a loss in global heterozygosity, and a need for frequent rescues might offset the benefit of restricting migration. It is also not clear that minimizing migration remains favorable under periodic rescues, even if the rate of rescues is independent of migration. To address these concerns, we derive a simple extension to the finite island model that incorporates the effects of hitchhiking during genetic rescues. We use the results of this model to argue that for a wide range of conditions, eliminating migration except as needed to allow genetic rescues will maximize global heterozygosity.

#### Core concepts and notation

We seek the dynamics of heterozygosity at a single genetic locus and follow the main text in defining heterozygosity as the probability that two gametes sampled with replacement from the given population carry different alleles. We denote heterozygosity with a lower-case $h$, and its expected value by the capitalized $H$. If $x_j$ is the frequency of the $j$-th allele in a population, then the heterozygosity of the population is given by $h = 1 - \sum_j x_j^2$, where the sum is over all possible alleles [1]. By this definition, $h$ depends only on allele frequencies, not genotype frequencies, and equals the fraction of heterozygous individuals only if the population is at Hardy-Weinberg equilibrium [1]. Given the initial state of the population at time $t = 0$, we wish to determine the heterozygosity at later times $t > 0$ (measured in generations). However, the allele frequencies that determine $h(t)$ evolve stochastically. Instead, we consider the expected value of the heterozygosity, $H(t) = E[h(t)]$, which approximates the value of $h(t)$ averaged over many independently evolving meta-populations or many independently evolving loci (in one meta-population) with the same initial conditions.

To find $H(t)$, we take advantage of its equivalent interpretation as the probability sampling different alleles at time $t$ knowing the state of the population at time 0. Equivalence follows from the following probability calculation: If $A$ is the event that 2 gametes sampled at $t$ are distinct alleles, then

$$\begin{aligned}
\Pr[A] &= \int_0^1 \Pr[A \mid h(t) = u] \Pr[h(t) = u]\, du \\
&= \int_0^1 u \Pr[h(t) = u]\, du \\
&= E\left[h(t)\right].
\end{aligned}$$

Interpreting $H(t) = E[h(t)]$ as the probability of sampling distinct alleles allows us to analyze its evolution without reference to the unknown underlying allele frequencies.

We seek the expected values of the *global heterozygosity* in the total meta-population and the *local heterozygosity*, or the average heterozygosity within local populations, which we hereafter refer to as *demes*. We use definitions of these quantities similar to Nei [3], modified to distinguish between the actual heterozygosity and its expected value (as noted above) and using the term "heterozygosity" instead of Nei's "gene diversity." Throughout, we consider a meta-population of $N_\mathrm{T} = LN$ total individuals subdivided into $L$ demes that contain $N$ diploid individuals. Conditional on the state of the meta-population at time $t$, we let $h_i(t)$ be the heterozygosity in deme $i$, $h_S(t)$ be the (average) local heterozygosity (with $h_S(t) = L^{-1} \sum_i h_i(t)$), and $h_T(t)$ be the heterozygosity the total population. Each $h_\bullet(t)$ equals the probability that two gametes randomly sampled (with replacement) at time $t$ are distinct alleles: $h_i$ corresponds to sampling from deme $i$, $h_S$ corresponds to sampling from a random deme, and $h_T$ corresponds to sampling from the total meta-population. We further define $h_D(t)$ as the probability that the gametes differ when sampled from two different demes. We note that $0 \leq h_S \leq h_T \leq h_D \leq 1$, with $h_T = h_S = h_D$ only when all demes have identical allele frequencies. We denote expectations of these quantities over the unknown meta-population state by $H_T(t)$, $H_S(t)$, and $H_D(t)$.

The relation $h_T = (1/L)h_S + (1 - 1/L)h_D$ can be found by conditioning on whether the second of two gametes sampled from the total meta-population are from the same or different demes. We use this relation often in its expectation form,

$$H_T = \frac{1}{L} H_S + \left(1 - \frac{1}{L}\right) H_D. \tag{1}$$

It will be useful to define a version of Wright's fixation index by $F_{\mathrm{ST}} = (H_T - H_S)/H_S$. There are many alternative, incompatible definitions of Wright's fixation index; we use this definition for its convenience in interpreting the following calculations. This quantity is not the same as the expected value of $(h_T - h_s)/h_T$ because in general $E[h_T/h_s] \neq E[h_T]/E[h_s]$, though the two may be approximately equal in many situations [4].

#### Finite island model: Drift, migration, and mutation

**Model:** First, we consider the finite-population version of Wright's island model [2, 5, 6], in which there are $L$ equally-sized demes. Each generation, each deme receives a fraction $m$ of migrants that are genetically representative of the total meta-population. For mathematical tractability and compatibility with traditional theory, we assume the locus to be selectively neutral. Each deme evolves according to the neutral Wright-Fisher model with mutation and migration, with migration of gametes rather than diploid individuals. The next generation is chosen by sampling with replacement from the previous generation to fill the $2NL$ available spots. With probability $1 - m$ sampling occurs from the same deme, and with probability $m$ from the total meta-population. (This definition of migration allows for migrants to end up in the same deme and is made for mathematical convenience. It amounts to a rate of migration to a different deme of $m(L - 1)/L$.) After each sampling, there is a probability $\mu$ of mutation, which gives rise to a new variant not yet in the meta-population. This so-called infinite alleles assumption is a good approximation when the locus consists of a long sequence of nucleotides [2]. Many details about the life cycle, such as whether migration occurs in gametes or individuals or whether generations are overlapping or discrete, are expected to have only minor effects on the results, provided the parameters are adjusted accordingly. In particular, a local effective population size $N_\mathrm{e}$ can be used in place of $N$ in the equations below to account for faster rates of drift than in Wright-Fisher sampling.

**Dynamics of expected total and local heterozygosity:** The effect of island-model subdivision on heterozygosity has been analyzed and described many times in the past [including by 1, 2, 6–9], but with various and sometimes subtle differences in assumptions and the specific quantities that are analyzed. To avoid the complexities of determining how previous analyses describe $H_S$ and $H_T$ in our model, we derive their dynamics anew. Thus the results we present below draw heavily from but are not directly equivalent to previous results.

To determine the expected heterozygosities through time, we first consider their change over a single generation. For compactness, let $H_S$, $H_D$, and $H_T$ be the values at time $t$ and $H'_S$, $H'_D$, and $H'_T$ be the values at time $t+1$. The expected local heterozygosity at time $t+1$ is

$$H'_S = \left(1 - \frac{1}{2N}\right)\left\{(1-\mu)^2\left([1-m]^2 H_S + \left[2m(1-m) + m^2\right] H_T\right) + 2\mu(1-\mu) + \mu^2\right\}. \quad (2)$$

This equation is computed by conditioning on the possible events pertaining to a sample of two gametes from a random deme at time $t+1$. There is a probability $1/2N$ that we sampled literally the same gamete (i.e., they are *identical by origin* [1]), in which case they are necessarily identical genetically (identical by state). Hence the leading factor $(1 - 1/2N)$ is the probability of sampling different gametes, and the term in curly braces is probability of sampling different alleles conditional on sampling different gametes. To determine this probability, we condition on whether either gamete experienced mutation. If so, they are necessarily distinct; if not, we further condition on whether either experienced migration. When either is a migrant, our sample traces back to a sample with replacement from the meta-population at time $t$, so that the gametes are distinct with probability $H_T$. If neither is a migrant, then our sample is equivalent to a sample with replacement from a random deme at time $t$ and thus distinct with probability $H_S$. Finally, we calculate the change $\Delta H_S = H'_S - H_S$ when the mutation and migration rates are small and the deme size $N$ is large. Rearranging (2) and ignoring terms that are second-order in $1/2N$, $\mu$, and $m$ yields

$$\Delta H_S \approx 2\mu(1 - H_S) + 2m(H_T - H_S) - \frac{1}{2N} H_S. \tag{3}$$

The three terms on the right-hand side of (3) describe the effects of mutation, which increases $H_S$ towards 1; migration, which increases $H_S$ towards the current value of $H_T$; and genetic drift, which decreases $H_S$ towards 0.

Repeating this procedure for the expected "different-deme" heterozygosity $H_D$, we obtain

$$H_D' = (1-\mu)^2 \left( [1-m]^2 H_D + \left[ 2m(1-m) + m^2 \right] H_T \right) + 2u(1-\mu) + \mu^2. \tag{4}$$

This equation now comes from conditioning on events pertaining to a sample of two gametes at $t + 1$ from two different demes. The term $(1 - 1/2N)$ is now gone, as the gametes are necessarily different by origin. Otherwise, the logic proceeds as for Equation (2). Rearranging and ignoring second order terms now leads to the approximate per-generation change,

$$\Delta H_D \approx 2\mu(1 - H_D) + 2m(H_T - H_D). \tag{5}$$

Mutation increases $H_D$ as with $H_S$, but the drift term from Equation (3) is gone, reflecting the lack of competition between gametes from different demes. As with $H_S$, migration drives $H_D$ towards the current value of $H_T$; however, since $H_D \geq H_T$, migration causes $H_D$ to decrease.

The expected total heterozygosity $H_T'$ is given by substituting (2) and (4) into (1). The change in expected total heterozygosity (again ignoring second-order terms) is similarly calculated from Equations (3) and (5), yielding

$$\Delta H_T \approx 2\mu(1 - H_T) - \frac{1}{2NL} H_S. \tag{6}$$

The effect of migration on $H_S$ and $H_D$ cancel, so that $H_T$ is unaffected by migration—as must be the case, given that migration does not change total allele frequencies—leaving just the effects of mutation and drift. The decrease in $H_T$ from drift is proportional to the reciprocal meta-population size, $1/NL$, and the local heterozygosity, $H_S$. This dependence on $H_S$ rather than $H_T$ arises because each gamete competes only with gametes from the same deme for a slot in the next generation.

Since the expected heterozygosities at $t+1$ only depend on their values at $t$, Equations (3) and (6) fully determine the trajectories $H_S$ and $H_T$ for all $t \geq 0$. We therefore use these equations in our numerical calculations. These equations also show that increasing the migration rate always has a short term effect of increasing $H_S$, which in turn accelerates the decay of $H_T$, while decreasing the migration rate always increases $H_T$.

**Minimal required migration to maintain local heterozygosity (Scheme 2):** In the absence of migration, the local heterozygosity will fall much more rapidly than the total heterozygosity. Under management Scheme 2, heterozygosity is monitored and the migration rate periodically adjusted to maintain $H_S$ above a critical threshold $c$. Here we consider how we might choose the new rate of migration. We suppose that, through genetic monitoring, we have estimates of both the average total and local heterozygosities. Using Equation (3) and ignoring the effect of mutation, we can see that maintaining $H_S$ at or above its current value requires $Nm \geq H_S/[4(H_T - H_S)]$. Substituting the threshold $c$ in for $H_S$ suggests we must chose a rate $Nm$ of migrants per deme per generation that satisfies $Nm \geq c/[4(H_T - c)]$. In practice we suggest picking $Nm$ to be somewhat larger than the minimum value to account for 1) we may only want to update the migration rate periodically, say every 10 or 100 generations; 2) the necessary migration rate increases as $H_T$ decreases; and 3) we can only imperfectly estimate the average heterozygosities. For instance, we might set

$$Nm = \frac{\alpha c}{4(H_T - c)}, \tag{7}$$

where $\alpha \geq 1$ is a parameter controlling how much larger than the current minimum to set the migration rate. This policy generally will involve less migration than a one-migrant-per-generation policy until the point where $H_T$ has also fallen to near the threshold. In particular, it sets $m = 0$ until $H_S$ reaches $c$, and afterward the new migration rate given by (7) will have $Nm < 1$ so long as $H_T > c(1 + \alpha/4)$.

### Genetic rescues

**Model:** Our model of genetic rescue is motivated by the following hypothetical scenario. After a focal deme is determined to be declining for genetic reasons (such as inbreeding depression or inadequate genetic variation to adapt to a changing environment), a small number of individuals from other demes are brought in to breed with the focal deme. If rescue is successful, one or more alleles at a non-neutral locus from the immigrant individuals will confer higher fitness on the offspring of the next generation. These beneficial allele(s) quickly increase in frequency in the rescued deme, causing linked alleles at nearby neutral loci to increase in frequency through the process known as genetic hitchhiking.

We approximate the effect of rescue on linked neutral diversity in this scenario with the pseudo-hitchhiking model [10]. The effect of a rescue is treated as instantaneous replacement of a fraction $R$ of gametes in the deme by the hitchhiking allele—the neutral allele that is linked to the sweeping beneficial variant. Approximating hitchhiking as instantaneous reflects our expectation that the sweep will be fast relative to the changes in heterozygosity by drift, mutation, and low rates of migration. For simplicity, we suppose that the hitchhiking allele is chosen randomly from the meta-population. We consider two scenarios for when rescues take place. In the first scenario, each deme is rescued with a constant probability of $\lambda$ per generation, with $\lambda \ll 1$. This scenario allows us to consider the long-term effects of periodic rescues on heterozygosity. In the second scenario, all demes are rescued simultaneously at a particular point in time. This scenario approximates Scheme 1 from the main text and is used to generate the simulated trajectory in Figure 2. The derivations above and the proposed management schemes can be numerically explored using the online computational tool at: https://ryantaylor.shinyapps.io/om2m/.

**Effect of rescues on total and local heterozygosity:** Because rescues occur instantly, we can consider their effect at a particular point in time in isolation from the other evolutionary processes of mutation, migration, and drift. We suppose that each deme experiences a rescue independently with probability $\lambda$ and compute the expected heterozygosities at time $t+1$ conditional on the meta-population at time $t$, which we denote $H'_S$, $H'_D$, and $H'_T$. As before, we compute these quantities by conditioning on the possible events that might befall our two sampled gametes, but now considering only events relating to genetic rescue.

By this procedure, we find that the expected local heterozygosity at $t+1$ is

$$H'_S = (1-\lambda)H_S + \lambda\left[(1-R)^2 H_S + 2R(1-R)H_T\right]. \tag{8}$$

Here we condition on whether the chosen deme experienced a rescue (with probability $\lambda$)

and if so, whether neither, one, or both of the sampled gametes was in the fraction $R$ of gametes derived from the hitchhiking allele. If no rescue occurred, or if a rescue occurred but neither gamete is in the hitchhiking fraction, then the gametes resemble a sample from time $t$ and the probability that they differ is $H_S$. If both gametes are in the hitchhiking fraction (probability $\lambda R^2$) then they are necessarily identical. If just one is, then they resemble a random sample from the meta-population at $t$ and hence differ with probability $H_T$. The change in $H_S$ equals

$$\Delta H_S = 2\lambda R(1-R)(H_T - H_S) - \lambda R^2 H_S. \tag{9}$$

Equation (9) shows that rescues have opposing effects on local heterozygosity. Comparing (9) to (3), we see that the first term of (9) represents an increase in $H_S$ that would occur with migration rate equal to $2\lambda R(1-R)$, and the second term represents a decrease in $H_S$ that would occur from drift with a local effective population size of $1/2\lambda R^2$.

The expected different-deme heterozygosity is

$$\begin{aligned} H'_D = (1-\lambda)^2 H_D + 2\lambda(1-\lambda)[(1-R)H_D + RH_T] \\ + \lambda^2[(1-R)^2 H_D + 2R(1-R)H_T + R^2 H_T]. \end{aligned} \tag{10}$$

This equation is derived similarly to (8), except the gametes are in different demes and so we now have to consider whether neither, one, or both demes experienced a rescue. If $\lambda \ll 1$, then we can ignore the $\lambda^2$ terms (reflecting the unlikely event that both demes were rescued) and the change in $H_D$ is approximately

$$\Delta H_D \approx 2\lambda R(H_T - H_D). \tag{11}$$

The scenario where all demes are rescued simultaneously is given by setting $\lambda = 1$; in this case, the change in $H_D$ is

$$\Delta H'_D = [2R(1-R) + R^2](H_T - H_D). \tag{12}$$

The expected total heterozygosity $H'_T$ is given by substituting (8) and (10) into (1), and the change in $H_T$ is found by similarly combining (9) and (11) or (12). When rescues are rare ($\lambda \ll 1$), the change in $H_T$ is approximately

$$\Delta H_T \approx -\frac{\lambda R^2}{L} H_S - \frac{2\lambda R^2}{L}(H_T - H_S). \tag{13}$$

The first term is from the drift-like decay term in Equation (9) for $\Delta H_S$. The second term is from the migration-like decrease in $H_D$, which is not entirely balanced from the migration-like increase in $H_S$ as it was with regular migration. When all demes are rescued simultaneously, the change in $H_T$ is

$$\Delta H_T = -\frac{R^2}{L} H_T. \tag{14}$$

This equation is used along with (9) (with $\lambda = 1$) for the simulation of Scheme 1 in the main text.

### Long-term dynamics

We now consider how subdivision, migration, and rescues affect the long-term dynamics of $H_T$. In particular, we follow the standard procedure of considering how the evolution of $H_T$ compares to its evolution in an unstructured population of equal total size [1, 2]. The change in $H_T$ in an unstructured population is given by setting $H_S = H_T$ in Equation (6), giving

$$\begin{aligned} \Delta H_T &\approx 2\mu(1 - H_T) - \frac{1}{2N_{\mathrm{T}}} H_T \\ &= 2\mu - \left(2\mu + \frac{1}{2N_{\mathrm{T}}}\right) H_T, \end{aligned} \tag{15}$$

which has the form

$$\Delta H = a - bH, \tag{16}$$

with $a = 2\mu$ and $b = -(2\mu + 1/2N_{\mathrm{T}})$. The solution of (16) is

$$\begin{aligned} H(t) &= \frac{a}{b} + \left(H(0) - \frac{a}{b}\right)(1 - b)^t \\ &\approx \frac{a}{b} + \left(H(0) - \frac{a}{b}\right) e^{-bt} \qquad \text{for } b \ll 1. \end{aligned} \tag{17}$$

That is, $H$ approaches an equilibrium of $\hat{H} = a/b$ exponentially at rate $b$. Thus in an unstructured population, the equilibrium is $\hat{H}_T = a/b = \theta/(1+\theta)$, where $\theta = 4N_{\mathrm{T}}\mu$, and is approached at rate $b = 2\mu + 1/2N_{\mathrm{T}}$ per generation.

To model the joint effects of subdivision, migration, and rescues, we can imagine that a generation is split into two phases: First, each deme is simultaneously and independently rescued with probability $\lambda$ (which we now assume is $\ll 1$); second, the meta-population experiences one generation of evolution under the finite island model. The changes in $H_S$ and $H_T$ are the sum of their respective changes over the two phases. When $\mu$, $m$, $1/2N$, and $\lambda$ are all small ($\ll 1$), the total change in $H_T$ can be approximated by summing Equations (6) and (13), giving

$$\Delta H_T \approx 2\mu(1 - H_T) - \frac{1}{2NL}H_S - \frac{\lambda R^2}{L}H_S - \frac{2\lambda R^2}{L}(H_T - H_S). \tag{18}$$

We can rewrite this equation in terms of $H_T$ and the fixation index $F_{\mathrm{ST}} = (H_T - H_S)/H_T$ to obtain

$$\Delta H_T \approx 2\mu(1 - H_T) - \left[\frac{1 - F_{\mathrm{ST}}}{2NL} + \frac{\lambda R^2}{L}(1 + F_{\mathrm{ST}})\right] H_T. \tag{19}$$

Similar equations appear in meta-population models (see below) with an assumed equilibrium value of $F_{\mathrm{ST}}$; here we make no equilibrium assumption, but simply use the algebraic relationships between $H_S$, $H_T$, and the present non-equilibrium definition of $F_{\mathrm{ST}}$. Equation (19) now has the same form as (15), but with the bracketed term replacing $1/2N_{\mathrm{T}}$. We can see that subdivision slows the decay in $H_T$ from genetic drift in proportion to $1 - F_{\mathrm{ST}}$. On the other hand, rescues increase the decay in $H_T$ through the additional term $\lambda R^2(1 + F_{\mathrm{ST}})/L$.

To determine how $H_T$ evolves in a subdivided population, we must still determine how $H_S$ (or $F_{\mathrm{ST}}$) evolves. Summing Equations (3) and (9) gives that the change in $H_S$ is approximately

$$\Delta H_S \approx 2\mu(1 - H_S) + 2m(H_T - H_S) - \frac{1}{2N}H_S + 2\lambda R(1 - R)(H_T - H_S) - \lambda R^2 H_S \tag{20}$$

which we rewrite as

$$\Delta H_S \approx 2\mu + 2(m + R\lambda - \lambda R^2)H_T - \left[2\mu + 2m + \frac{1}{2N} + \lambda R(2 - R)\right] H_S. \tag{21}$$

If $H_T$ were fixed, this equation would have the form of (16) and $H_S$ would tend to an equilibrium of

$$\hat{H}_S \mid H_T = \frac{4N\mu + 4N[m + \lambda R - \lambda R^2]H_T}{1 + 2N[2\mu + 2m + 2\lambda R - \lambda R^2]}. \tag{22}$$

Comparing the bracketed term in (21) to that in (19) shows that if the number of demes is large ($L \gg 1$), then $H_S$ approaches $\hat{H}_S \mid H_T$ much faster than $H_T$ approaches its equilibrium. As a result, $H_S$ will approximately reach $\hat{H}_S \mid H_T$ and track this value as $H_T$ changes, allowing us to determine the subsequent dynamics of $H_T$ by assuming $H_S = \hat{H}_S \mid H_T$. The quantity $\hat{H}_S \mid H_T$ is often referred to as a quasi-equilibrium [8], and such quasi-equilibrium approximations when $L \gg 1$ are frequently used to analyze the long-term dynamics of meta-populations [8, 11–14].

If we ignore the effect of new mutations on $H_S$ by setting $\mu = 0$, then the ratio of $H_S/H_T$ at quasi-equilibrium is independent of $H_T$. As a result, $F_{\text{ST}}$ has a true equilibrium of

$$\hat{F}_{\text{ST}} \approx \frac{1 + 2N\lambda R^2}{1 + 4Nm + 2N\lambda R(2 - R)}. \tag{23}$$

Substituting $\hat{F}_{\text{ST}}$ for $F_{\text{ST}}$ in Equation (19) leads to

$$\Delta H_T \approx 2\mu(1 - H_T) - \left[\frac{1 - \hat{F}_{\text{ST}}}{2NL} + \frac{\lambda R^2}{L}(1 + \hat{F}_{\text{ST}})\right] H_T. \tag{24}$$

Comparing this expression to (15) for an unstructured population, we find the effective size of the population with recurrent rescue events to be

$$N_{\text{eT}} \approx \frac{NL}{1 - \hat{F}_{\text{ST}} + 2N\lambda R^2(1 + \hat{F}_{\text{ST}})}. \tag{25}$$

Equations (23) and (25) are similar to expressions for $\hat{F}_{\text{ST}}$ and $N_{\text{eT}}$ in other meta-population models; see Whitlock [15] and our discussion of extinction-recolonization models below.

In the absence of rescues, we have Wright's classic result [15]

$$N_{\text{eT}} \approx \frac{N_{\text{T}}}{1 - \hat{F}_{\text{ST}}} = N_{\text{T}} \cdot \frac{4Nm}{1 + 4Nm}, \tag{26}$$

which makes clear that decreasing migration always increases $N_{\text{eT}}$ and thus total heterozygosity in the absence of rescues.

When does decreasing $m$ help maintain global heterozygosity in the presence of recurrent rescues and when all other parameters are fixed? Equations (23) and (24) show that migration affects $H_T$ indirectly through $\hat{F}_{\mathrm{ST}}$. Decreasing the migration rate always increases $\hat{F}_{\mathrm{ST}}$. Meanwhile, greater values of $\hat{F}_{\mathrm{ST}}$ decrease the decay in $H_T$ due to drift (through the term $(1-\hat{F}_{\mathrm{ST}})/2NL$) and increase the decay in $H_T$ due to rescues (through the term $\frac{\lambda R^2}{L}(1+\hat{F}_{\mathrm{ST}})$). Decreasing the migration rate therefore favors the maintenance of $H_T$ when $1/2N > \lambda R^2$, so that the decay in $H_T$ is mainly from drift rather than rescues. Since the migration rate $m$ is absent from the condition, the optimal strategy for maintenance of global heterozygosity is to have maximal migration if $1/2N < \lambda R^2$ and no migration if $1/2N > \lambda R^2$.

An important caveat is that this condition does not account for the possibility that lower levels of local heterozygosity increase the required rate of rescues. The long-term consequences of such an effect regarding policy choices for migration are far from clear, since a low migration policy can maintain higher levels of $H_S$ in the mid to long term despite giving lower $H_S$ in the short term (main text Figure 2). The effects of various migration policies when rescues (or local extinctions, considered below) depend on local population genetics therefore warrant further exploration.

### Comparison with extinction-recolonization models

Our model can be seen as an extension of previous models designed to capture the effects of local extinction-recolonization dynamics on heterozygosity in populations [9, 12, 16–18]. Comparing the results of our rescue model with these previous results provides insight into the generality of the condition that determines when decreasing migration rates preserves total heterozygosity. It also clarifies some seemingly contradictory statements about the effects of subdivision on heterozygosity in these models.

These models of extinction-recolonization dynamics [9, 12, 16–18] assume island model dynamics with demes additionally going extinct with a constant probability $\lambda$ per generation. Demes experiencing an extinction are instantly repopulated by a set of founder gametes or diploid individuals. When only one founder gamete is chosen, these models resemble our rescue model with $R = 1$. Following the notation of Whitlock and Barton [9] and Roze and Rousset [12], let the number of founders equal $k$ individuals or $2k$ gametes, with $k = 1/2$ corresponding to a single gamete. How the founders are chosen is determined by a parameter $\phi$, the probability that founder gametes come from the same deme, with $\phi = 0$ corresponding to a random sample from the total population and $\phi = 1$ corresponding to gametes coming from just one deme. Models differ in whether founders are chosen before or after the extinction event, which determines whether it is possible for founder alleles to come from the deme experiencing extinction; however, resulting differences in expressions for heterozygosity and $N_{\mathrm{eT}}$ are minor if $\lambda \ll 1$.

To verify our results and assess their generality, we compare our expressions for $\hat{F}_{\mathrm{ST}}$ and $N_{\mathrm{eT}}$ and the condition for when decreasing the migration rate preserves total heterozygosity to those from three relatively recent studies of extinction-recolonization models. Cherry [18] model a haploid population in which the founder allele is a random sample from the total population before the extinction event. Their approximations for $\hat{F}_{\mathrm{ST}}$ (unnumbered equation at the top of their p. 790) and $N_{\mathrm{eT}}$ (their Equation 11) exactly agree with our equations (23) and (25) after adjusting for diploidy. Whitlock and Barton [9] and Roze and Rousset [12] consider a diploid population with general $k$ and $\phi$. In their models, founder alleles are chosen from the remaining demes after any extinctions. They find the following approximation for $\hat{F}_{\mathrm{ST}}$

$$F_{\mathrm{ST}} \approx \frac{1 + \frac{N\lambda}{k}}{1 + 4Nm + 2N\lambda\left[1 - \phi\left(1 - \frac{1}{2k}\right)\right]};$$

see Whitlock and Barton [9] Equation (21) and Roze and Rousset [12] Equation (34). This expression reduces to ours (with $R = 1$) when $k = 1/2$. For the effective population size with $\phi = 0$, Whitlock and Barton [9] Equation (22) gives

$$N_{\mathrm{eT}} = \frac{NL(1-\lambda)}{1 - \hat{F}_{\mathrm{ST}} + 2N\lambda\hat{F}_{\mathrm{ST}}}, \tag{27}$$

which differs from our expression and that of Cherry [18] in the denominator. Although this expression differs from ours, it leads to a consistent condition for decreasing migration to favor total heterozygosity. The derivative of $1/2N_{\mathrm{eT}}$ with respect to migration rate is positive if and only if

$$\frac{1}{2N} > \lambda\left[1 + \phi\left(1 - \frac{1}{2k}\right)\right]. \tag{28}$$

This condition reduces to our condition (with $R = 1$) when $k = 1/2$ or $\phi = 0$. Rousset

[19] point out that Equation (27) is inconsistent with the assumptions of the model used by Whitlock and Barton [9] (originally derived from Slatkin [16]), but that the approximation in Whitlock and Barton [9] Equation (23),

$$N_{\mathrm{eT}} \approx \frac{L}{4(m+\lambda)\hat{F}_{\mathrm{ST}}}. \tag{29}$$

is valid for small $\lambda$ and $m$. This approximate form for $N_{\mathrm{eT}}$ also leads to the inequality (28). This more general condition shows that migration is more likely to be beneficial when $k$ and $\phi$ are larger, but that $1/2N > 2\lambda$ is sufficient to favor reducing migration regardless of how founder alleles are chosen.

This sufficient condition under the more general extinction model has important implications for our proposed policy. It suggests that $1/2N > 2\lambda$ is also a sufficient condition for low to no migration to be preferred in a conservation setting when $\lambda$ is the combined rate of extinctions and rescues, regardless of how demes are rescued or repopulated. An important caveat as noted above is that this condition does not account for the possibility that lower levels of local heterozygosity increase the rate of rescues and extinctions, an issue that requires further exploration.

We conclude our discussion of extinction models by examining claims that subdivision decreases $N_{\mathrm{eT}}$ and total heterozygosity. For example, the abstract of Whitlock and Barton [9] concludes: "Contrary to the expectation from the standard island model, the usual effect of population subdivision is to decrease the effective size relative to a panmictic population living on the same resource." This statement appears to suggest that increasing the degree of subdivision by decreasing the migration rate between demes would typically reduce $N_{\mathrm{eT}}$, rather than increase it. However, a careful reading of this study shows that the reductions in $N_{\mathrm{eT}}$ being referred to are due to local extinctions, source-sink dynamics, or other ways in which demes vary in their genetic contributions to subsequent generations. This pattern appears to hold true across other studies of extinction-recolonization dynamics [16, 17]. These studies demonstrate that increasing the extinction rate reduces $N_{\mathrm{eT}}$ and draw attention to the fact that, unlike in the standard island model, the effective size $N_{\mathrm{eT}}$ can be lower than the census size $NL$. They do not, however, consider the effect of changing the migration rate $m$ on $N_{\mathrm{eT}}$. To our knowledge, we are the first to consider the question of when a decrease of the migration rate increases total heterozygosity in the presence of factors, such as rescues or extinction-recolonization dynamics, that may cause $N_{\mathrm{eT}}$ to be lower than the census size $NL$. We find that $1/2N > 2\lambda$ is a sufficient condition for $N_{\mathrm{eT}}$ to be maximized by minimizing migration, and there is a broad range of conditions under which "subdivision" broadly construed might be interpreted as reducing $N_{\mathrm{eT}}$ below $NL$ and yet decrease in $m$ would increase $N_{\mathrm{eT}}$.

**Supplementary 3: Review and discussion of related studies and test cases**

**Contents:**

**3.1 Inbreeding**
**3.2 Insight regarding population dynamics from case-studies of small populations**
**3.3 Genetic rescue**
**3.4 What do studies and management plans recommend with regard to connectivity?**

**Supplementary 3.1: inbreeding**

Although a thorough review of the literature on inbreeding, bottlenecks, and genetic rescue is well beyond the scope of the current study, we highlight a number of important observations.

***Supplementary 3.1 highlights***

- For species conservation, inbreeding is a risk only if it leads to population-level decline.
- The observation of fitness reduction that is limited to highly inbred individuals is of secondary importance for species conservation. Such an observation may suggest the occurrence of effective purging of deleterious alleles, and may be interpreted as a positive signal of the population's viability.
- It is well established that inbreeding significantly reduces fitness at the individual level; this has been shown to also occur in the wild.
- It is often assumed that there is a linear relation between the inbreeding coefficient and fitness. This seems not to be the case in many studied species; instead there appears in some cases to be a threshold effect, where only extremely highly inbred individuals exhibit reduced fitness.
- Few studies have explored inbreeding effects at the population level, and findings are mixed.
- A number of case-studies suggest that some highly inbred populations are viable over evolutionary timescales, and there are well-studied cases in which highly successful populations originated from a handful of individuals.
- Recent studies have shown that genetic rescue of highly inbred populations may act rapidly and effectively to increase mean fitness and lead to population-level growth.
- Outbreeding depression seems to have limited effects, or does not occur at all, in many of the cases studied thus far, including cases of hybridization between populations that are considered separate subspecies.

Long-standing interest in inbreeding stems in part from its well-documented consequences in husbandry of domesticated species (reviewed in, e.g., [1–4]). Despite this effort, understanding of inbreeding remains limited and the subject is controversial. For example, a number of mechanisms have been proposed to underlie inbreeding effects, and their relative roles remain somewhat debated [3,4]: partial dominance (increased expression of deleterious recessive alleles), over-dominance (heterozygote advantage), and epistasis (increased likelihood of preferable allele combinations among heterozygotes). The first mechanism, partial dominance, is most often considered the most prevalent [5]. A second aspect that remains largely unresolved is how and how much inbreeding interacts with environmental factors: some studies propose that inbreeding depression is magnified in stressful or variable environments [6], an observation that might be interpreted as supporting a role for epistasis in bringing about inbreeding depression. A third topic of broad debate is the extent, rate, and efficacy of purging of deleterious alleles in highly inbred populations [3,4,7–9].

These issues have important implications in attempting to avoid population-level inbreeding depression. A particularly important question in this respect is the functional relation between the inbreeding coefficient and the observed fitness: is it a linear relationship, as is often assumed, or is it non-linear, with the effects of inbreeding being disproportionally large for highly-inbred individuals? The answer depends on a combination of the mechanisms mentioned above, as well as additional factors such as the mode of selection, population dynamics, and the species' life history. One such factor is whether selection is *hard*, eliminating members from the population, for example, every individual below a threshold body size, or whether it is *soft selection*, eliminating from the population, for example, a certain fraction of individuals every year, with the smallest individuals being the most likely to be removed, regardless of their absolute size. Some life-history traits may have similar influence, for example: if the mating system only allows the largest males reproduce, it may lead to a functional form in the relation between inbreeding coefficient and fitness that is different from that arising if reproduction were limited by an external factor, independent of the size of other individuals in the population, and all individuals above a certain body size managed to reproduce. These factors are rarely considered.

Many studies have shown inbreeding depression in wild populations (see, e.g., [4,10–13]). However, relating these findings to species' conservation and management of fragmented populations is not straightforward: the vast majority of these studies did not study whether inbreeding depression significantly affects population-level dynamics, and – most importantly – whether it may lead to population decline (but see, e.g. [14,15]). The fact that individuals with higher inbreeding coefficients fare worse than others does not imply that the population is at increased risk of extinction or that it is more vulnerable than other populations. Moreover, the *opposite* might be true: reduced fitness of highly-inbred individuals suggests that purging of deleterious recessive alleles may occur, decreasing the deleterious load in the population (suggested in, e.g. [16,17]). For example, inbreeding depression is clearly seen in humans: genetic disease, including lethal diseases, are common in offspring of close relatives, but humans are not in any way at risk of extinction due to inbreeding. This might not be true, of course, if all individuals in the population were highly inbred.

Although the population-level dynamics that are associated with inbreeding are not easily predictable (because they may interact, as noted, with the species' life history, ecology, specific environment, and population-specific phenomena such as competition) individual-based studies can be used to inform expectations regarding the possible influence of inbreeding at the population level. It would seem reasonable to assume that as long as many individuals in the population are not highly inbred, the population is not at increased risk due to inbreeding. When most individuals are highly inbred, this may not be the case. What constitutes a high inbreeding coefficient is an empirical question, and if data on inbreeding's effect on the species of interest is available, and the distribution of inbreeding coefficients in the population is known, predictions can be made regarding population dynamics.

We would like to highlight in this respect the importance of the functional relation between the inbreeding coefficient and fitness: if the function is non-linear, and fitness decreases suddenly above a certain threshold in the value of the inbreeding coefficient, it would suggest that management plans should aim to avoid a situation in which most of the individuals in the population have inbreeding coefficients near or above that threshold. If the functional form is linear and there is no natural threshold, a direct population-level measure, such as a measurement of population stability or decline, seems to be a useful criterion for when intervention is necessary to mitigate extinction risk. When possible, the two approaches to determining management aims and policy can be used in unison, i.e. avoiding a certain independently-inferred level of inbreeding, while also monitoring population-level growth or decline.

Most studies assume a linear relationship, based on simplistic assumptions, namely that (1) inbreeding is due to low per-individual heterozygosity and partial or over-dominance at many sites, and that (2) fitness is a linear function of additive contributions from each genetic site that influences it and at which the individual may be homozygous or heterozygous (see discussion of the topic in, e.g., [18–21]). We find the first assumption reasonable [5], but the latter is, in many cases, unlikely to be correct: at a minimum, it is incorrect whenever there is a disproportional reproductive skew among individuals with respect to the trait of interest (e.g. if reproduction is limited by available territories or nesting sites, and body size differences, for example, disproportionately influence the probability of territory acquisition). Many other individual-level mechanisms and population-level dynamics may invalidate the second assumption, for example between-locus epistatic effects on the phenotype and selection dynamics on reaching adulthood that do not respond linearly to trait values. The extent to which deviation from the second assumption influences the functional relation between inbreeding coefficient and fitness is unknown.

The relation between the inbreeding coefficient and fitness is an empirical one, and can be inferred from data; review of the studies that include such an analysis or that provide a visualization of the distribution of individuals' fitness scores as a function of their inbreeding coefficients yields mixed results. Some fitness proxies, such as first-year survival in red deer [12], seem to be in linear correlation with the inbreeding coefficient, while calves' weight at birth, in the same study, is characterized by a threshold function, and only individuals inbred at a level that is analogous to half-siblings' mating suffer a statistically-significant reduction in fitness. Similarly, the effect of heterozygosity on parasite load in populations of Galapagos hawks [22], the effect of parental relatedness on reproductive success in grey seals, pilot whales, and wandering albatrosses [23], and the overwinter survivorship of Soay sheep lambs [16] seem to follow a linear relationship, while the relation between measures of inbreeding and egg hatching probability in great reed warblers [24] and in blue tits [25], overwinter survivorship of adult Soay sheep [16], and first year survival of arctic foxes [26] seem to follow a threshold function. Unfortunately, many studies provide only summary statistics regarding inbreeding's effect on fitness, making it difficult to assess the extent to which one form or the other is true or whether one may generalize about which fitness proxies tend to follow which functional form.

A more direct way to address the question of interest – does inbreeding lead to population-level decline – is the study of whole populations or proxies that may correlate with the population's wellbeing. Reed and Frankham [27] conducted an influential meta-analysis of such studies. They were able to find only 34 such studies, among which only 5 were in vertebrates. They found that overall, inbreeding (used in this supplementary section as shorthand for decreased expected heterozygosity) may in many cases influence population-level measures of wellbeing, which are expected to correlate with risk of extinction. We suggest that these findings should be interpreted with caution. First, since only 5 of these studies were of vertebrates, the generalizability of findings from other taxa than vertebrates is unclear. Among the 5 studies, only 3 showed a significant influence of inbreeding on population-level measures of fitness, and 2 found no correlation. Second, in all 3 studies that reported a correlation between inbreeding and fitness, the prominent driver of the correlation was the data from a single population whose inbreeding coefficient was exceptionally high. Third, in all three studies, none of the populations – including those that were highly inbred – seem to be in decline; the inbred populations in all three studies presumably existed in isolation for hundreds or thousands of generations. In one of these studies, the population density even increased with the decrease in heterozygosity, with the greatest density found in the most inbred population. Finally, we note that although the authors are aware of the potential confounding effect of reporting bias, in which studies that found no significant correlation between inbreeding and population-level dynamics are less likely to be published, they could not effectively account for the extent to which this might occur.

To summarize, we do not assert that population-level decline never occurs as a result of inbreeding (see also [4]). We suggest that given the existing data, the jury should still be out regarding this question, and that the evidence is far from providing a decisive answer to it. Moreover, we point out that the mere fact that no sweeping evidence has been found in favor or in refutation of this relationship despite the great research efforts on this topic, suggests that the effect of inbreeding on population-level dynamics is either highly variable among populations, or of a limited size in general. Accordingly, we suggest that among conservation scientists, the common perception of the risk to populations due to inbreeding is out of proportion given the limited extent to which this risk is substantiated by studies of the topic.

**Supplementary 3.2: Insight regarding population dynamics from case-studies of small populations**

Many fragmented populations are small, and are likely to remain so in the foreseeable future. Studies that explored the demographic dynamics and the genetics in populations under similar conditions are useful for deriving insight about what we might expect in recently-fragmented populations under various conditions and alternative management plans. There are two prominent categories of such case studies: island populations, many of which have been separated from other populations for long periods of time, and reintroduced populations or native populations that reached an extremely low population size.

Island populations are "natural experiments" that demonstrate the fate over long timescales of fragmented populations in the absence or near-absence of between-population migration. The management paradigm that we propose in this study suggests maintenance of such conditions as the default management strategy, and that between-population migration be facilitated only when necessary, in the form of attempting genetic rescue if a population shows genetics-related decline or has reached a state in which most individuals in the population are inbred beyond a certain threshold that is deemed critical. Hence, island populations are a promising setting from which to make predictions about the proposed paradigm. The most important finding from studies of island populations (which is so obvious that it sometimes goes unnoticed) is the mere fact that these populations exist. Moreover, in many cases they seem to have endured for thousands of years, with typically small or intermediate population sizes and, quite likely, despite recurring bottlenecks (e.g. [28–32])[1]. Many island populations are characterized by low heterozygosity (e.g. [16,28,33–36]) and some studies of island populations have also found reduction in the mean absolute fitness of island populations, according to various proxies, in comparison to their more heterozygous mainland counterparts [28,34].

The endurance of these populations over thousands of years, suggests that populations may be less sensitive than is frequently assumed to reduction in their absolute fitness (even at the whole-population level) compared to an "optimal" genotype of their species. It may be, for example, that the limiting factor that determines the survival of a population is the availability of appropriate habitat, and as long as it exists, a reduction of fitness, even a significant one, has

[1] We include also studies of mainland populations that had been disconnected from other populations for exceedingly long time periods.

little influence on the probability of the population's survival. Thus, for example, it seems that black-footed rock wallabies on Barrow island suffer from a decrease in their reproductive success compared to their mainland counterparts; however this may not have mattered for the population's survival, which seems to have been at the island's carrying capacity for this species until recent anthropogenic threats were introduced [28,37]. Interestingly, in deer mice ([33]), despite a 12-fold higher parasite load among the individuals that inhabit Whiskey island compared to the parasite load in mainland populations (presumably related to the formers' high inbreeding coefficient), the population on the island exhibited the highest density among the studied islands. The same pattern was seen in this study across all island populations: the higher the inbreeding coefficients, the higher the parasite load and the population density. These increased densities may, at least in part, explain the increased parasite load independently of inbreeding.

Concerning management of small populations from a genetic perspective, the question of interest is whether inbreeding tends to influence the eventual extinction of island populations or not. This is a different problem, and has not been addressed, to the best of our knowledge. One can easily imagine, for example, that eventual extinction of island populations is driven by punctuated ecological challenges of large effect size, such as the sudden loss of the species' ecological niche on the island. Such a change would lead to the extinction of the population even if it had the absolute fitness that characterizes its mainland counterpart. Interestingly, a related argument has been made in the past, suggesting that the process leading to extinction occurs so rapidly, that the populations' genetic diversity is not significantly reduced before the extinction occurs (see [38–40]). On the other hand, it has been shown that some populations do in fact go through a stage in which their genetic diversity is reduced before they go extinct (e.g. [32,41]). However, this does not show that genetics had any significant influence on the extinction process (as is discussed in [42]).

An important point to keep in mind is that the sample of populations studied is severely biased: we can study primarily those that survived, while many others existed but went extinct. Thus these populations' long term survival despite reduced genetic diversity is encouraging, but it provides a demonstration of a population's possible trajectory, not of its probability. Assessing this probability requires knowing how many original populations existed compared to those that survived. This may be possible by analyzing a combination of genetic and biogeographic data (considering probabilities of re-colonization and other factors), but is complex, would yield results that are largely species- and context-specific, and well beyond the scope of the current study. Thinking in these terms, however, may be useful: for recently-fragmented populations, the current situation can be thought of as somewhat analogous to simultaneous colonization of multiple islands, that then remained disconnected from the mainland and largely also from one another.

In this “thought experiment”, from a genetic perspective, two important factors may have played a role in determining which of the original island populations survived. The first is the founder effect: the stochastic initial genetic composition of each population may have strongly influenced its survival, and in particular, island populations that initially happened to contain fewer lethal or near-lethal recessive alleles may have survived with higher probability. The second factor is purging: populations that survived might be the ones in which purging of deleterious alleles took place successfully, perhaps during particularly severe bottlenecks.

Following the above, a valid concern is that drawing insight from island populations to recently-fragmented populations may be misleading, since island populations may differ in their genetic characteristics from recently-fragmented populations. However, we suggest that to some extent, many extant fragmented populations might be characterized by founder effects and purging: fragmentation was recent, occurring primarily over the recent 200 years. Compared to typical time scales of genetic dynamics of neutral or near-neutral diversity, this is a short period; but for some purging and some influence of lethal alleles among the founding populations, processes that involve strong selection on small populations, this time period may have been sufficient. In other words, in many fragmented species, it may be that a few decades have already passed since the populations were fragmented, and the fragmented populations that still exist are more likely to be the ones that happened to avoid particularly problematic founding effects or to purge some alleles of large deleterious effect-size. If this is the case, the existing populations are more similar to island populations than one would expect from random population fragments: they are a subsample of all original fragments, with a bias towards population-level genetic compositions that are particularly viable and happen to have a reduced genetic load, such that inbreeding depression is not too severe.

Exploring case studies of species’ reintroduction from small pools of founders, or cases in which a native species was reduced to a very small population while being studied, can aid in providing perspectives that differ from the study of island populations with regard to the possible caveats discussed above. Unlike island populations, these bottlenecked populations do not have a long history in a state of disconnection, and they may represent a less biased sample in respect to the possible genetic-related fates of such small populations. Some well-studied such cases are cheetahs [43,44], Asiatic wild ass [45–47], and elephant seals [48]. In many of these cases, despite the population's reduction to few individuals, it increased in size dramatically, once appropriate habitat was available and direct (demographic) risk factors, particularly poaching, were removed. Although the bottleneck phase still influences the genetics of these populations, which are all still characterized by small effective population sizes, their populations seem highly successful and their effective population sizes are on the rise. In some cases, this increase is in part a result of informed management of the population or meta-population, which considers both demographic and genetic factors. Other populations in similar settings are, or were, less successful [49,50]. In these cases the population failed, or is still struggling, to increase in size following the bottleneck. Whether this is due to genetic factors is unknown in all of these cases, but given the small population sizes – significantly smaller than those that we find advisable for a management unit (for avoiding fixation of deleterious alleles of large effect; see main text and supplementary 4), genetics should not be ruled out, at least as a contributing factor. In the paradigm we propose, unless a population is in decline for reasons that are clearly non-genetic (e.g. poaching or frequent road kill), genetic rescue from another population should be facilitated. Insights from studied cases of genetic rescue are discussed in the following subsection, and its role and implementation in the paradigm we propose are discussed in supplementary 2.

We do not assert that island populations have endured at small population sizes and low heterozygosity for eons and that these factors do not affect risk of extinction, nor that the handful of cases of successful reintroductions from small pools of founders suggest that population size is unimportant. Obviously, the decreased fitness that may be associated with low heterozygosity is undesirable. However, as we show in the main text and in supplementary sections 3 and 4, increasing the heterozygosity via migration comes at a cost of decreasing the global diversity and with it, an increased probability of extinction due to environmental challenges that the lost alleles could potentially have mitigated. The studied cases of populations overcoming severe bottlenecks or enduring on islands for hundreds of generations, often with reduced genetic diversity, and latent or observed decrease in fitness, show that populations may survive and be successful with mean fitness that is significantly reduced compared to the species' optimum (e.g. [34]). Keeping this in mind may be useful in constructing management plans that balance the two costs.

**Supplementary 3.3: Genetic rescue**

Here, *Genetic rescue* is used to describe a scenario in which the introduction of a limited number of migrants into a population increases the population's absolute fitness and the population growth via an increase in genetic diversity [51–53]. This may occur because the migrants' contribution to the gene pool alleviates widespread inbreeding depression or because it introduces genetic adaptations to ecological challenges. The concept of *assisted gene flow* is similar, and highlights the case in which the environmental conditions that the receiving population faces changed in recent times; migrants under this scheme are introduced from a population in which the environmental conditions are more similar to the current conditions in the receiving population [51]. *Evolutionary rescue* partially overlaps our use of the term genetic rescue, and refers specifically to the case where a population declined as a result of an environmental change, followed by the spread of a genetic adaptation that allows the population to meet this challenge, leading to population growth. The source of the adaptation can be de-novo mutation, standing variation, or – in the case that it overlaps with genetic rescue – migrants from another population [53,54].

Genetic rescue as a result of facilitated migration of few individuals into populations that had been in decline has been reported in many studies, including in wild populations reviewed, for example, in [51]). As one might expect, the most significant effects occur when the migrants originated in large and diverse populations; however, genetic rescue is effective even when the source of migrants is itself highly inbred. Crossing between Drosophila lines, each of which had been inbred for multiple generations, leads to rapid genetic rescue, with fitness benefits seen in the first generation following the introduction of migrants and lasting for many generations thereafter [55–59]. The reported cases in which the population trajectory switched from decline to growth following genetic rescue include vertebrates [60], such as the Florida Panther [61], Gray wolf [62,63], Mexican wolf [64,65], Bighorn sheep [66–68], Wood rat [69], and European Viper [70]. In some of these cases it is hard to gauge the relative importance for the shift in population dynamics of the genetic rescue compared to the importance of other conservation measures such as habitat reconstruction and reduction in exposure to parasites. However, the emerging conclusion from these cases is that genetic rescue may have large effects on population-level dynamics, even on a short time scale and following migration of few individuals.

There are many considerations that must be taken into account in implementing genetic rescue via facilitated migration, including the species' life history, which may dictate which migrants might be most likely to induce genetic rescue (e.g. males or females, juveniles or adults), the risk of spreading disease between populations, and the risk of outbreeding depression [71]. Accumulating studies, summarized in a number of reviews on the topic [51,53,54,65], suggest useful guidelines to accommodate these considerations and minimize the risk involved. Apart from the risk of reducing overall genetic diversity, which is the reason that we advocate the implementation of genetic rescue only when necessary, outbreeding depression seems to be the risk factor that is most difficult to control. However, it seems that there is negligible risk of outbreeding depression when rescue is attempted between populations that diverged recently, and empirical evidence suggests that even admixture of populations that diverged so early as to constitute different subspecies, i.e. whose common ancestor existed tens of thousands of years ago, may be highly beneficial and should be attempted in cases where a population is severely inbred[2] or declining and alternative sources for genetic rescue are unavailable.

[2] As discussed in the section about inbreeding, empirical data may suggest a clear population-level inbreeding threshold that should be avoided, below which individual fitness drops precipitously. In the absence of such information, we suggest that genetic rescue be attempted if a population reaches a state in which most individuals are homozygous to the extent expected in mating between siblings or half-siblings.

**Supplementary 3.4: What do studies and management plans recommend with regard to connectivity?**

Most recovery plans for terrestrial vertebrates stress the "genetic health" of the population(s) or species being considered, and many recommend gene-flow as a prescription for poor genetic health. Rarely do studies differentiate between within-population genetic diversity and between-population genetic diversity (but see, e.g., [72,73]). Inbreeding depression is considered by many a primary determinant of extinction risk of fragmented populations, and strong recommendations regarding the necessity of increasing between-fragment connectivity are often made in the study or management plan of fragmented species (e.g. [74–78]). However, as discussed above, extremely few studies have shown population-level decline due to inbreeding. In particular, we note that even when empirical findings explicitly suggest that there is no indication of inbreeding depression at the population level, researchers often recommend increasing population connectivity, either via ecological corridors and animal crossings across barriers or via facilitated migration of individuals. For example, Miller et al. [79] studied the genetics of grizzly bears in Yellowstone and found that diversity is low, but that it has changed little over the last century. They do not observe reduced values in proxies for fitness, and conclude that inbreeding depression does not pose a challenge to this population; however, the authors still recommend increasing connectivity between this population and others. The 2012 jaguar recovery outline [77], prepared for US Fish and Wildlife, recommends establishing corridors between jaguar populations despite there being no evidence of inbreeding in any of the populations, and mixed evidence of genetic differentiation. The 2007 bighorn sheep recovery plan posits that "Stemming further loss of genetic diversity in Sierra Nevada bighorn sheep may prove to be the greatest challenge for their recovery", and recommends the maintenance of gene flow across all populations (despite acknowledgement of the potential importance of global species diversity). Notably, in many cases the phrasing of recommendations makes it clear that the authors believe increased connectivity supports local diversity as well as global diversity, not considering that often – particularly in the setting of a population that crashed in recent decades or centuries – there is a tradeoff between the two, and increase in one is at the expense of the other (e.g. [75,77]).

wallaby in the Warburton region of Western Australia. *Wildl. Res.* **19**, 605–621.

38. Lande, R. 1988 Genetics and demography in biological conservation. *Science (80-. ).* **241**, 1455–1460.
39. Caro, T. M. & Laurenson, M. K. 1994 Ecological and genetic factors in conservation: a cautionary tale. *Sci. Pap. Ed. Guid. to Sci. Inf.* **263**, 485–486.
40. Elgar, M. A. & Clode, D. 2001 Inbreeding and extinction in island populations: a cautionary note. *Conserv. Biol.* **15**, 284–286.
41. Spielman, D., Brook, B. W. & Frankham, R. 2004 Most species are not driven to extinction before genetic factors impact them. *Proc. Natl. Acad. Sci. U. S. A.* **101**, 15261–15264.
42. Jamieson, I. G. & Allendorf, F. W. 2012 How does the 50/500 rule apply to MVPs? *Trends Ecol. Evol.* **27**, 578–584.
43. Dobrynin, P. et al. 2015 Genomic legacy of the African cheetah, Acinonyx jubatus. *Genome Biol.* **16**, 277.
44. Merola, M. 1994 A reassessment of homozygosity and the case for inbreeding depression in the cheetah, Acinonyx jubatus: implications for conservation. *Conserv. Biol.* **8**, 961–971.
45. Gerard, J.-F., Ziv, A., Bouskila, A. & Bar-David, S. 2015 Space-use patterns of the asiatic wild ass (Equus hemionus): complementary insights from displacement, recursion movement and habitat selection analyses. *Plos One 12*
46. Renan, S., Greenbaum, G., Shahar, N., Templeton, A. R., Bouskila, A. & Bar-David, S. 2015 Stochastic modelling of shifts in allele frequencies reveals a strongly polygynous mating system in the re-introduced Asiatic wild ass. *Mol. Ecol.* **24**, 1433–1446.
47. Gueta, T., Templeton, A. R. & Bar-David, S. 2014 Development of genetic structure in a heterogeneous landscape over a short time frame: the reintroduced Asiatic wild ass. *Conserv. Genet.* **15**, 1231–1242.
48. Weber, D., Stewart, B. S., Garza, J. C. & Lehman, N. 2000 An empirical genetic assessment of the severity of the northern elephant seal population bottleneck. *Curr. Biol.* **10**, 1287–1290.
49. Seddon, P. J., Armstrong, D. P. & Maloney, R. F. 2007 Developing the science of reintroduction biology. *Conserv. Biol.* **21**, 303–312.
50. Armstrong, D. P. & Seddon, P. J. 2008 Directions in reintroduction biology. *Trends Ecol. Evol.* **23**, 20–25.
51. Whiteley, A. R., Fitzpatrick, S. W., Funk, W. C. & Tallmon, D. A. 2015 Genetic rescue to the rescue. *Trends Ecol. Evol.* **30**, 42–49. (doi:https://doi.org/10.1016/j.tree.2014.10.009)
52. Tallmon, D. A., Luikart, G. & Waples, R. S. 2004 The alluring simplicity and complex reality of genetic rescue. *Trends Ecol. Evol.* **19**, 489–496.
53. Carlson, S. M., Cunningham, C. J. & Westley, P. A. H. 2014 Evolutionary rescue in a changing world. *Trends Ecol. Evol.* **29**, 521–530.
54. Gonzalez, A., Ronce, O., Ferriere, R. & Hochberg, M. E. 2013 Evolutionary rescue: an emerging focus at the intersection between ecology and evolution.
55. Bijlsma, R., Westerhof, M. D. D., Roekx, L. P. & Pen, I. 2010 Dynamics of genetic rescue in inbred Drosophila melanogaster populations. *Conserv. Genet.* **11**, 449–462.
56. Spielman, D. & Frankham, R. 1992 Modeling problems in conservation genetics using captive Drosophila populations: improvement of reproductive fitness due to immigration of one individual into small partially inbred populations. *Zoo Biol.* **11**, 343–351.

57. Ball, S. J., Adams, M., Possingham, H. P. & Keller, M. A. 2000 The genetic contribution of single male immigrants to small, inbred populations: a laboratory study using Drosophila melanogaster. *Heredity (Edinb).* **84**, 677–684.
58. Ávila, V., Fernández, J., Quesada, H. & Caballero, A. 2011 An experimental evaluation with Drosophila melanogaster of a novel dynamic system for the management of subdivided populations in conservation programs. *Heredity (Edinb).* **106**, 765.
59. Åkesson, M., Liberg, O., Sand, H., Wabakken, P., Bensch, S. & Flagstad, Ø. 2016 Genetic rescue in a severely inbred wolf population. *Mol. Ecol.* **25**, 4745–4756.
60. Trinkel, M. et al. 2008 Translocating lions into an inbred lion population in the Hluhluwe‐iMfolozi Park, South Africa. *Anim. Conserv.* **11**, 138–143.
61. Johnson, W. E. et al. 2010 Genetic restoration of the Florida panther. *Science (80-. ).* **329**, 1641–1645.
62. Vilà, C., Sundqvist, A., Flagstad, Ø., Seddon, J., Kojola, I., Casulli, A., Sand, H., Wabakken, P. & Ellegren, H. 2003 Rescue of a severely bottlenecked wolf (Canis lupus) population by a single immigrant. *Proc. R. Soc. London B Biol. Sci.* **270**, 91–97.
63. Adams, J. R., Vucetich, L. M., Hedrick, P. W., Peterson, R. O. & Vucetich, J. A. 2011 Genomic sweep and potential genetic rescue during limiting environmental conditions in an isolated wolf population. In *Proc. R. Soc. B*, pp. 3336–3344. The Royal Society.
64. Fredrickson, R. J., Siminski, P., Woolf, M. & Hedrick, P. W. 2007 Genetic rescue and inbreeding depression in Mexican wolves. *Proc. R. Soc. London B Biol. Sci.* **274**, 2365–2371.
65. Hedrick, P. W. & Fredrickson, R. 2010 Genetic rescue guidelines with examples from Mexican wolves and Florida panthers. *Conserv Genet* **11**, 615–626.
66. Hogg, J. T., Forbes, S. H., Steele, B. M. & Luikart, G. 2006 Genetic rescue of an insular population of large mammals. *Proc. R. Soc. London B Biol. Sci.* **273**, 1491–1499.
67. Miller, J. M., Poissant, J., Hogg, J. T. & Coltman, D. W. 2012 Genomic consequences of genetic rescue in an insular population of bighorn sheep (Ovis canadensis). *Mol. Ecol.* **21**, 1583–1596.
68. Poirier, M., Coltman, D. W., Pelletier, F., Jorgenson, J. & Festa‐Bianchet, M. 2018 Genetic decline, restoration and rescue of an isolated ungulate population. *Evol. Appl.*
69. Smyser, T. J., Johnson, S. A., Page, L. K., Hudson, C. M. & Rhodes, O. E. 2013 Use of experimental translocations of Allegheny woodrat to decipher causal agents of decline. *Conserv. Biol.* **27**, 752–762.
70. Madsen, T., Shine, R., Olsson, M. & Wittzell, H. 1999 Conservation biology: restoration of an inbred adder population. *Nature* **402**, 34.
71. Waller, D. M. 2015 Genetic rescue: a safe or risky bet? *Mol. Ecol.* **24**, 2595–2597.
72. Service, U. S. F. and W. 2013 *Recovery Plan for the Black-footed Ferret (Mustela Nigripes).* Region 6, US Fish and Wildlife Service.
73. Service, U. S. F. and W. 2007 *Recovery plan for the Sierra Nevada bighorn sheep.* California/Nevada Operations Office, US Fish and Wildlife Service.
74. Coleman, R. A., Weeks, A. R. & Hoffmann, A. A. 2013 Balancing genetic uniqueness and genetic variation in determining conservation and translocation strategies: a comprehensive case study of threatened dwarf galaxias, Galaxiella pusilla (Mack)(Pisces: Galaxiidae). *Mol. Ecol.* **22**, 1820–1835.
75. Yuasa, T., Nagata, J., Hamasaki, S., Tsuruga, H. & Furubayashi, K. 2007 The impact of habitat fragmentation on genetic structure of the Japanese sika deer (Cervus nippon) in

southern Kantoh, revealed by mitochondrial D-loop sequences. *Ecol. Res.* **22**, 97–106.

76. Vonholdt, B. M., Stahler, D. R., Bangs, E. E., Smith, D. W., Jimenez, M. D., Mack, C. M., Niemeyer, C. C., Pollinger, J. P. & Wayne, R. K. 2010 A novel assessment of population structure and gene flow in grey wolf populations of the Northern Rocky Mountains of the United States. *Mol. Ecol.* **19**, 4412–4427.
77. Team, J. R. & Service, U. S. F. and W. 2013 Recovery Outline for the Jaguar (Panthera onca).
78. Haag, T. et al. 2010 The effect of habitat fragmentation on the genetic structure of a top predator: loss of diversity and high differentiation among remnant populations of Atlantic Forest jaguars (Panthera onca). *Mol. Ecol.* **19**, 4906–4921.
79. Miller, C. R. & Waits, L. P. 2003 The history of effective population size and genetic diversity in the Yellowstone grizzly (Ursus arctos): implications for conservation. *Proc. Natl. Acad. Sci.* **100**, 4334–4339.

#### Supplementary 4: A brief summary of population structure theory

Population genetic theory includes many models in which genetic variation and evolution in structured populations are studied. These models vary in their underlying assumptions and the evolutionary process they study. Much of the literature has been concerned with measuring the consequences of population subdivision for local and global genetic variation measured by F-statistics [1]. The variance in allele frequencies due to population structure can be measured as $F_{st}$, namely the proportion of the total variance in allele frequencies contained in subpopulations relative to the total allele-frequency variance. $F_{st}$ is, therefore, also a measure of genetic differentiation among the subpopulations.

For the Infinite Island Model, where migration is assumed to be identical between all demes at a fixed rate $m$ (measured as the proportion of the population migrating each generation), and all demes are assumed to be of the same effective size $N$, it can be shown that, approximately, at equilibrium [1]:

$$F_{ST} = \frac{1}{1+4Nm} (S4.1)$$

Eq. S4.1 shows that, under the model assumptions, the balance between local and global genetic variation depends on the absolute number of migrants per generation, $Nm$, rather than the relative rates. Observing from Equation S4.1 that for $Nm = 1$ we expect 80% of the total variance to be contained within demes ($Fst = 0.2$), it has been suggested as a general guideline for the purpose of conservation that migration at a rate of one migrant per generation (OMPG) between demes would keep much of the variance at the local level [2,3]. This would prevent low levels of heterozygosity within local populations and allow meta-populations to avoid the negative consequences of inbreeding depression [4]. It would also mean that differentiation between demes is not too large, and subpopulations would be kept relatively genetically similar.

The effect of population subdivision can also be observed from the perspective of its impact on effective population size. The effective population size is the size of an ideal population (randomly mating, fully admixed population) that experiences genetic drift at a similar rate to the population being studied, with respect to a given genetic measure [5,6]. For example, the variance effective population size, $N_{ev}$, is the size of an ideal randomly mating population that experiences accumulation of allele-frequency variance at a similar rate to our focal population; the inbreeding effective size, $N_{ef}$ is similarly defined with respect to the accumulation of inbreeding, typically measured by the probability that two alleles at a given locus in an individual are identical by descent (coefficient of inbreeding). The consequences of population structure on effective sizes have been explored for example in the Finite Island Model. For example, a finite island model with a meta-population of size $nN$ is partitioned to $n$ demes of size $N$, and the variance effective size for the total population is [7]:

$$N_{ev} = \frac{nN}{1 - F_{ST}} \, (S4.2)$$

Using Eq. S4.1 to describe $F_{st}$, we have:

$$N_{ev} = nN\left(1 + \frac{1}{4Nm}\right) (S4.3)$$

This indicates that with population structure, low absolute migration rates result in higher variance effective population sizes (higher than the actual population size), i.e. the more the population is structured, the lower is the global rate of accumulation of allele frequency variance through time.

$N_{ef}$ shows an opposite trend, and under similar assumptions with a mutation rate of $\mu$, we have for the total inbreeding effective size [8]:

$$N_{ef} = N\left(1 + \frac{m}{\mu}\right) (S4.4)$$

and, globally, inbreeding is greater with more structuring.

Some models have described population structure in a more spatially explicit manner, such as the stepping-stone models [9] and the continuous isolation-by distance-model [10]. These models add another level of complexity with long- and short-distance migration rates. However, the overall conclusion that increased population structuring (decreased migration) results in higher level of global genetic variation, is maintained.

Another class of models considers meta-population dynamics and the possibility that demes might go extinct, and later be colonized by other demes. Such dynamics are not discussed in the main text, as we consider scenarios where subpopulations can be properly maintained and are no longer at direct demographic risk. Typically, in models where extinctions and colonizations are allowed, the variance effective size decreases with population subdivision, contrary to the scenario described above [11]. This is because when an extinction-colonization event occurs, the meta-population loses the diversity found in one deme, and instead now has two "copies" of the same deme. This increases genetic homogeneity in the meta-population and decreases genetic diversity globally. Consequently, in meta-population models with extinction/colonization, the balance between the extinction/colonization rates and between-deme differentiation determines whether population structure increases or decreases variance effective size and rates of loss of genetic diversity [11].

The models described above mostly address neutral genetic variation and the balance between migration and genetic drift. Other models have attempted to analyze the balance of migration, drift and selection under the 'shifting balance theory' [1,12]. For example, Barton & Rohani [13] found that relatively low or moderate absolute migration rates, of the order of $Nm = 1$ or even lower, may allow meta-populations to advance more efficiently towards a global adaptative optimum through consecutive shifts through the 'adaptive landscape' than occurs with higher migration rates, i.e. more highly connected meta-populations. Similarly, taking into account epistasis (interactions of genes at different loci), Hadany [14] showed that some low migration rates allow the most efficient shifting-balance type evolution, while with high connectivity evolution is hindered.

*Genetic diversity*

Different aspects of genetic diversity are measured in different ways. The most common measure of genetic diversity in conservation studies is heterozygosity, either observed heterozygosity (proportion of heterozygous individuals in the population) or expected heterozygosity , which we consider in the current study ($1 - \sum_i p_i{}^2$, where $p_i$ are the allele frequencies in the population). These measures are particularly related to inbreeding, as mating between individuals with similar genetic makeup (e.g. relatives) results in individuals with higher homozygosity than expected. Low heterozygosity is therefore often regarded as an indicator of inbreeding depression (see *supplementary 1*).

Genetic diversity can also be measured by allelic diversity, the number of different alleles in the population. Allelic diversity is an indicator of adaptive potential, since different alleles are the raw material upon which selection may act—the more alleles present in a population the higher the chance that one of these alleles will be beneficial under some environmental conditions, and consequently driven to fixation by natural selection. This is particularly important in cases where environmental conditions are expected to change (e.g. global warming), since alleles that are currently under no or low selection pressure, may in the future be targets of selection. The theory regarding allelic diversity in fragmented populations is far less developed then that of heterozygosity, partly due to the fact that allelic diversity models are less mathematically tractable than those involving heterozygosity (see [15–18]).

*The conservation genetic literature*

The conservation literature has been very much interested in the evolutionary consequences of fragmented populations, as these have become more common with anthropogenic interference. Whereas most models in the general population genetic literature describe the balance of local and global genetic variation and their evolutionary consequences as a result of the interaction of migration and genetic drift, most models in the conservation literature are aimed at providing guidelines for management of populations, with some genetic goal in mind.

In particular, Eq. S4.1 and the OMPG have been used to argue that, with inbreeding in local population as a negative consequence to be avoided, exchange of *at least* one migrant per generation between demes should be maintained (e.g. [4,19,20]). The robustness of this threshold to violations of the assumptions used to derive Eq. S4.1, with the same genetic goal of avoidance of deleterious local inbreeding effects, has also been investigated [4,21,22]. However, it is not often emphasized that the OMPG rule assumes that the populations are at migration-drift equilibrium, an assumption that is often incorrect in conservation scenarios, where populations are going, or have recently undergone, an extensive reduction in size. Therefore, even when considering the goal of the OMPG rule (to avoid local inbreeding depression), it is only when the population reaches the new equilibrium that this rule should be applied, and migration should be scaled to the actual heterozygosity level until then.

The goal of avoiding local inbreeding depression has been the main concern in many theoretical conservation studies. Several have investigated extinction risk as a result of inbreeding in non-fragmented populations [23,24], while some have addressed fragmented populations in detail with computer simulations of specific scenarios to estimate the threshold above which inbreeding depression does not occur. For example Kenney et al. [25] modeled inbreeding depression in tigers under different migration rates to assess various management strategies.

Fewer studies in conservation biology address the issue of maintenance of evolutionary potential by keeping global allelic diversity at high levels [17]. This neglects the importance of allelic diversity for the long-term survival of populations – allelic diversity is a better indicator of adaptive potential than heterozygosity [16,26,27]. Fewer studies still ask how different migration schemes affect allelic diversity retention. Greenbaum et al. [16] estimated allelic diversity retention in theoretical conservation scenarios with different migration rates under a Continent-Island model, and Weiser et al. [28] simulated specific conservation scenarios for several species. Caballero et al. [17] simulated both heterozygosity and allelic diversity under different migration regimes for the critically endangered *Turdus helleri* to evaluate the most useful strategy. The message from these studies is that migration regimes aimed at avoiding low heterozygosity and inbreeding at the within-deme level are not necessarily the best regimes for maximizing allelic diversity and evolutionary potential [16,17].

Many population genetic and conservation genetic models have been aimed at understanding the consequences of population structure for fragmented populations. However, most focus on inbreeding-depression avoidance, and pay less attention to the effect of migration on global genetic diversity. Moreover, these models fail to incorporate an important characteristic of currently endangered populations, namely that they often still retain much of the genetic diversity of their ancestral large population. This diversity is being eroded as the populations advance towards the new small-size equilibria. Therefore, we encourage studies that consider the dynamics that are expected to play out until the new equilibria are reached, and that acknowledge that assisted migration, while beneficial for preventing inbreeding, has costs for global genetic diversity of the meta-population.

**Supplementary 5: Supporting calculation for the illustrative example in the main text**

We consider a species whose population, say, 200 years ago, was at size $N_0 = 1{,}000{,}000$ individuals (effective size), and was reduced to $N_I$ = 1000 individuals (effective size), split between 10 populations. We assume that the original population had been fully connected for a long time and was at Hardy-Weinberg equilibrium. With these conditions, under the neutral theory [1], and assuming the infinite-sites mutation model [2] (appropriate for loci spanning at least several SNPs, in which homoplasy is unlikely), the expected heterozygosity is predicted to be ([3], p. 112):

$$H = \frac{\theta}{1+\theta} (S5.1)$$

with $\theta = 4N\mu$. For the purpose of this example, we are considering a neutral locus with mutation rate $\mu = 10^{-5}$. The expected heterozygosity of the ancestral population is therefore $H_0 = 0.97$. The expected allelic diversity ($A$) at equilibrium can be calculated by considering the allele frequency spectrum [4]:

$$A = \int_{\frac{1}{2N_0}}^{1} \frac{(1-x)^{\theta-1}}{x} dx \ (S5.2).$$

With the parameters above, we have at time zero $A_0$ =410.2.

Now we consider a subsample of size $N_1 = 1{,}000$ of this large population. For simplicity, we will assume an instantaneous reduction in size. The effect of subsampling on heterozygosity is similar to a single generation of genetic drift in an unstructured population of size $N_1$, which does not change heterozygosity significantly, leaving $H_1 = 0.97$. Subsampling of allelic diversity in this manner has been addressed by Ewens ([4] and [3], p.114), and the expected allelic diversity after such sampling would approximately be:

$$A = \theta \int_0^1 \frac{(1-x)^{\theta-1}}{x} (1-(1-x)^{2N_1}) \approx \frac{\theta}{\theta} + \frac{\theta}{\theta+1} + \cdots + \frac{\theta}{\theta+2N_1-1} (S5.3)$$

We can now obtain the expected allelic diversity in our example immediately after the population reduction, $A_1 = 157.8$.

Next, we suppose that in recent times, the species was fragmented into $K = 10$ populations, and consider the implications of different management schemes for the equilibrium genetic diversity. In particular, we contrast the extreme cases of maximizing and minimizing migration between populations. For the case where the population is managed as a single panmictic population, the long-term equilibrium heterozygosity can be calculated using Eq. S5.1 with the new meta-population size $N_1 = 1{,}000$, giving $H_2 = 0.04$. The equilibrium allelic diversity is calculated using Eq. S5.2, and we obtain the equilibrium allelic diversity, $A_2 = 1.3$.

Next we consider that the population is kept fragmented to $K$ populations with no migration between them. For simplicity, we ignore new mutations in our calculations of total heterozygosity and allelic richness, which provides a conservative lower bound for their ultimate values. In this case, one allele is fixed in each population at equilibrium, thus neglecting the added diversity contributed by mutation (mutation-drift equilibrium). In each deme, the probability of any allele to fix is the same as the frequency of the allele in the deme at the onset of the genetic process [5], i.e. immediately after the population size reduction. Therefore, in order to determine which allele type would be brought to fixation in each deme we must sample a single gene from the initial deme gene pool. Since the deme gene pool itself is a subsample of the entire meta-population after size-reduction, and this gene pool is in turn a subsample of the pre-reduction meta-population gene pool, the allele that will ultimately be fixed can be determined by a single gene sample from the large population. We have K demes, so in order to determine the allelic diversity at fixation equilibrium we must consider the sampling of K genes from the original population, a problem identical to the one addressed by Ewens [4], and we have the estimate, similarly to Eq. S5.2:

$$A \approx \frac{\theta}{\theta} + \frac{\theta}{\theta + 1} + \cdots + \frac{\theta}{\theta + K - 1} \quad (S5.3)$$

With our example, $K = 10$, and thus the fixation equilibrium of allelic diversity, which represents a lower bound on the migration-drift equilibrium of allelic diversity, is $A_2 = 9.0$. The equilibrium expected heterozygosity of the meta-population can be found by noting that two gametes sampled now at random from the meta-population will be in the same population with probability 1/K and in different populations with probability (K-1)/K. In addition, two gametes sampled from the same population will be identical, and two gametes from different populations are equivalent to two gametes sampled from the initial population (in terms of their lineages' coalescence). Thus, ignoring new mutations, the equilibrium expected heterozygosity is

$$H_2 = \frac{K-1}{K} \cdot H_1 = 0.87 .$$

Without mutation, the within-population heterozygosity would drop to 0, which is unrealistically low. Instead, we note that its value with mutation can be calculated from Eq. S5.1 with N = N1 / 10 = 100. Within-population heterozygosity is thus expected to drop to the extremely low value of approximately 0.004.

To summarize, under the simplistic assumptions of this toy example, high genetic diversity was found in the original large population, and much of it is still represented after the crash in population size and the fragmentation ($H_0 = H_1 = 0.97$; $A_0 = 410.2$ and $A_1 = 157.8$). If managed as a single highly-connected population, most genetic diversity will be lost, and at equilibrium the species would reach $H_2 = 0.04$ and $A_2 = 1.3$. If maintained as completely separate populations, the within-population diversity would be very low, but globally high genetic diversity would be retained: $H_2 = 0.87$ and $A_2 = 9.0$ (here $A_2$ is the fixation equilibrium, a lower bound for the mutation-drift equilibrium).
